# Having the ‘right’ microbiome matters for host trait expression and the strength of mutualism between duckweeds and microbes

**DOI:** 10.1101/2022.02.10.479958

**Authors:** Anna M. O’Brien, Jason Laurich, Megan E. Frederickson

**Affiliations:** Dept. of Ecology and Evolutionary Biology, University of Toronto

**Keywords:** duckweed, species interactions, adaptation, GxG, GxE, urban pollution

## Abstract

An organism’s phenotypes and fitness often depend on interactive effects of its genome (*G*_*host*_), microbiome (*G*_*microbe*_), and environment (*E*). These *G* x*G, G* x*E*, and *G* x*G* x*E* effects fundamentally shape host-microbiome (co)evolution and may be widespread, but are rarely compared within a single experiment. We collected and cultured *Lemna minor* (duckweed) and its associated microbiome from 10 sites across an urban-to-rural ecotone. We factorially manipulated host genotype and microbiome in two environments (low and high zinc, an urban aquatic stressor) in an experiment with 200 treatments: 10 host genotypes *×* 10 microbiomes *×* 2 environments. Host genotype explained the most variation in *L. minor* fitness and traits, while microbiome effects depended on host genotype (*G* x*G*) or environment (*G* x*E*). Hosts had higher fitness and microbes grew fastest when tested microbiomes more closely matched field microbiomes, suggesting some local adaptation between hosts and their microbiota. High microbiome similarity also led to more predictable host trait expression. In contrast, although zinc decreased host fitness, we observed no local adaptation of urban duckweed or microbes to high-zinc conditions. Thus, we found that host fitness and trait expression are contingent on microbiome composition, with implications for microbiome engineering and host-microbiome evolution.

## Introduction

The microbial communities making a living on larger hosts are an integral part of their host’s ecology and evolution, affecting trait expression and fitness in hosts across the tree of life (Honegger, 1993; Turnbaugh et al., 2008; Friesen et al., 2011; Gould et al., 2018; Dittami et al., 2016). In turn, host trait variation can affect microbial survival, growth, or transmission (Adler et al., 2021; Buffington et al., 2016). Theoretical models suggest that microbiomes will contain genetic variation for host traits and that this variation will respond to selection on hosts (Rebolleda-Gómez et al., 2019; O’Brien et al., 2021; Henry et al., 2021). Emerging empirical results support this idea: altering microbiomes inoculated onto hosts can change the additive genetic variance and covariance of traits (O’Brien et al., 2019), and both host traits and host fitness can ‘evolve’ in microbiomes when host genotypes are held constant (Lau and Lennon, 2012; Panke-Buisse et al., 2015; Tso et al., 2018; Batstone et al., 2020). These and similar results have spurred continued calls to engineer microbiomes for agricultural, medical, or other applications (e.g. Epstein et al., 2019; French et al., 2021; Khan et al., 2021).

We expect that much ‘evolution’ of host traits in microbial genomes or communities results from links between microbial and host fitness via reciprocal fitness feedbacks (Sachs et al., 2004). For example, in plants hosting nitrogen-fixing microbes, microbe-derived nitrogen may allow the plant host to grow larger, which in turn may translate into more plant-fixed sugars provided to microbes, and thus a positive fitness feedback between partners (Friesen, 2012). Variation in host fitness caused by expressed host trait values could therefore impact the relative fitness of microbial genotypes or species that have variable effects on those traits. Indeed ‘evolved’ effects of microbes on host phenotypes have been linked to both community composition turnover and evolution within strains (Lau and Lennon, 2012; Panke-Buisse et al., 2015; Tso et al., 2018; Batstone et al., 2020). Furthermore, when hosts and microbes share an evolutionary history, ongoing fitness feedbacks may cause local adaptation between them (Rúa et al., 2016; Houwenhuyse et al., 2021).

The dependence of expressed trait values and heritable variation for traits on microbiome context is not unique to the microbiome environment, however. Context-dependent effects of genotypic variation underlie local adaptation to contrasting physical environments (Anderson et al., 2013), a common and well-known evolutionary phenomenon (Hereford, 2009). Like microbiomes, physical environments can simultaneously change trait expression, the amount of heritable genetic variation, and selection pressures (Wood and Brodie III, 2015, 2016). Furthermore, physical environments may alter the evolutionary forces that act on the variation for host traits and fitness contained in microbiomes (Bertness and Callaway, 1994; Bronstein, 1994; O’Brien et al., 2018). For example, nutrient pollution disrupts positive fitness feedbacks between hosts and microbial symbionts (Shantz et al., 2016), and evolution in microbes in response to nutrient loading can lead to fitness reductions in hosts (Weese et al., 2015).

Before setting out to engineer or evolve host traits in microbiomes, we must ask how much trait variation actually depends on the microbiome versus host genotype, and whether microbiome effects are predictable or contingent on host genotype or physical environment. Yet, estimating microbial effects on host traits is rarely conducted across host genotypes and in multiple environments, as it requires high-throughput experiments (but see O’Brien et al., 2019; Zahn and Amend, 2019; Fitzpatrick et al., 2019; Chaney and Baucom, 2020). The common duckweed *Lemna minor* and its largely mutualistic, manipulable microbiome (O’Brien et al., 2020) is a useful model for such experiments because of *L. minor* ‘s small size and fast generation time. Here, we carried out a full-factorial *G × G × E* experiment using duckweed and microbes to directly compare the relative importance of host genotype (*G*_*host*_), microbiomes (*G*_*microbe*_), physical environment (*E*), and their interactive effects (*G*_*host*_ *× G*_*microbe*_, *G*_*host*_ *× E, G*_*microbe*_ *× E*, and *G*_*host*_ *× G*_*microbe*_ *× E*) to host phenotype and fitness.

Duckweed is a floating aquatic plant with a wide distribution across diverse environments (RDSC, 2016), including both rural and urban sites. Urban-to-rural gradients can be stark ecotones, and may drive a substantial amount of evolution (Johnson and Munshi-South, 2017): perhaps most famously in soot-matching peppered moths (Kettlewell, 1955), but more recently in freshwater organisms dealing with increasingly toxic stormwater inputs from dense urban zones (Kern and Langerhans, 2018; Brans and De Meester, 2018). From previous work, we know that duckweed-associated microbes can alleviate some, but not all, urban stressors (O’Brien et al., 2019), and fail to ameliorate the effects of zinc (O’Brien et al., 2020), which duckweed can take up from water (Dirilgen and Inel, 1994; Radić et al., 2010; Jayasri and Suthindhiran, 2017). Zinc is a common urban runoff contaminant because it occurs in car tires and building roofs (Göbel et al., 2007), and duckweed exposure to zinc therefore varies along the urban-to-rural ecotone (Liskco and Struger, 1996; Göbel et al., 2007; Ontario Ministry of the Environment, 2011).

We tested all possible combinations of 10 clonal *L. minor* lines and the culturable portion of their field-collected microbiomes in two environments: high and low zinc. We expected to find that both host genotype and microbiome contribute to variation in duckweed fitness and traits, perhaps contingent on contamination context. We further hypothesized that duckweeds and microbes benefit from shared ecological and evolutionary history, and from an ecological history of zinc exposure in high-zinc conditions: i.e. that local adaptation underlies some *G*_*host*_ *× G*_*microbe*_, *G*_*host*_ *× E*, and *G*_*microbe*_ *× E* effects. More generally, *G*_*host*_ *× G*_*microbiome*_ effects may be detectable only when the difference in microbiomes is sufficiently large, and more diverged microbiomes may be more likely to produce non-additive effects, in much the same way that more diverged genomes cause more epistasis (Dettman et al., 2007; De Visser et al., 2011). Therefore, we also predicted that more dissimilar microbiomes are more likely to affect duckweed hosts in genotype-specific ways: i.e. that *G*_*host*_ *× G*_*microbiome*_ effects will be larger when comparing more dissimilar microbial communities.

## Methods

### Biological materials

In May-August 2017, we collected common duckweed, *Lemna minor*, and its associated microbes from 10 sites in the Greater Toronto Area (see Table S1), spanning a gradient from urban to suburban to rural. From each site, we isolated microbes from 1-2 duckweed fronds by pulverizing freshly collected plant tissue and streaking onto an agar plate with yeast-mannitol (YMA) media, which we incubated at 29^*°*^C for 5 days before storing at 4^*°*^C until use in the experiment. These microbes thus comprise the subset of *L. minor* epiphytic and endophytic microbes that can be maintained in mixed culture on YMA media. We also collected approximately 50 additional *L. minor* fronds from each site which we froze for 16S rRNA gene profiling of field *L. minor* microbiomes. Finally, we isolated one living *L. minor* frond from each site. We maintained the progeny resulting from individual field fronds in growth media (Krazčič et al., 1995, refreshed every *≈*2 weeks), in vented glass 500 mL mason jars, in a growth chamber (ENCONAIR AC80, Winnipeg, Canada) with a cycle of 16 hours at 23^*°*^C at 150 *µ*mol/m^2^ and 8 hours of 18^*°*^C in the dark. Fronds grew from a single individual to hundreds in about three months, and were maintained at high density until the experiment.

In the field, *L. minor* produces new fronds mainly clonally (i.e., by budding), but can flower and reproduce sexually via seed. However, genetic evidence shows that this occurs rarely (we never observed flowers in our cultures) and that there is little segregating diversity within populations, meaning that even sexually produced isoparental offspring may be nearly isogenic (Ho, 2017). Therefore, our *L. minor* cultures derived from a single field-collected fronds are effectively isogenic.

### Sequencing of microbial communities

To characterize and compare the microbiomes of field-collected *L. minor* and the inocula we used in experiments, we used 16S rRNA amplicon sequencing. We extracted DNA from field collected frozen duckweed tissue with DNeasy Powersoil kits (Qiagen, Hilden, Germany), using the recommended fresh tissue weight (0.25 g). We extracted DNA from our cultured microbiomes with GenElute Bacterial Genomic DNA kits (Millipore-Sigma, St. Louis, MO, USA). We sent *≈*10 ng of DNA from each extraction to Genome Qúebec (McGill University) for PCR amplification of the 16s rRNA gene (V3-V4 region, 341f/805r, Table S2, good sequence recovery and little bias expected for bacteria in plant microbiomes, Thijs et al., 2017), normalization, and barcoded, paired-end, 250 base pair format sequencing on an Illumina MiSeq System (San Diego, CA, USA). We received *≈*962 million base pairs in *≈*1.9 million demultiplexed reads, ranging from 26,029 to 129,507 reads across samples.

We used QIIME2 software to process reads. We first trimmed adapters and primers, joined paired reads, and quality filtered, including trimming to 402 bp, the length at which quality declined (identified with QIIME2’s visualization tools). We then assigned amplicon sequence variants (ASVs) and corrected sequencing errors with tool ‘deblur,’ and inferred taxonomic affiliation of ASVs using the naive Bayesian ‘classify-sklearn’ method trained on the Greengenes database (McDonald et al., 2012). We limited recording taxonomy identifications to those inferred with a confidence of 70% or higher, and we filtered out reads identified as chloroplast (Streptophyta only) or mitochrondria. We inserted reads into the Greengenes reference phylogeny (Janssen et al., 2018), discarding those that could not be placed. After all processing steps, 514,747 reads remained, ranging from 9,430 to 49,135 across samples (mean 25,737). Field samples had a similar number of reads per sample as cultured samples pre-processing, but fewer post-processing (average 15,826 for field samples versus 36,349 for cultured samples post-processing). Field samples were extracted from plant tissue, and contained numerous Streptophyta chloroplast reads (average of 16,560 per sample), accounting for the post-processing difference in depth.

We used QIIME2’s gneiss tool to compute the contrast in relative abundance of taxa subtending each daughter branch from selected nodes of the tree (‘balances,’ Morton et al., 2017). The selected nodes were those where sequences from any daughter ASV of that node were found in at least 8 of the 10 cultured communities. We also constructed pairwise distance matrices of each sequenced microbial community to the other using abundance weighted Unifrac distance (Lozupone and Knight, 2005, hereafter ‘Unifrac distance’), and plotted distances and taxonomic breakdown and ASV overlap across paired field and cultured community samples in R (R Core Team, 2019).

### Experimental design and data collection

We performed fully factorial manipulation of duckweed from 10 sites, duckweed microbial communities from those same sites, and the presence or absence of an environmental stressor: high concentration of zinc.

We set the high zinc treatment to 3.44 *µ*M (approximate level in runoff from an urban highway or waste discharge, Liskco and Struger, 1996; Göbel et al., 2007; Ontario Ministry of the Environment, 2011), by adding ZnSO_4_ to growth media. We used the unaltered media for the low zinc treatment, in which zinc was present at 0.86 *µ*M because it is a micronutrient. Zinc is also naturally present at low levels in duckweed habitat (Liskco and Struger, 1996). To prepare microbial inocula, we stirred a swab from the stored agar plate for each site into separate tubes of liquid YMA, which we then cultured in a shaking incubator (VWR, Radnor, PA, USA) for 3 days at 30 ^*°*^C and 200 rpm. We diluted these using a calibration curve for optical density to cell density in order to end up with 2,000 cells/*µ*L. Our optical density transformation is predictive of colony forming units in mixed communities (O’Brien et al., 2020), but it is imperfect: the relationship between optical density and number of live cells can vary both across and within microbial species (Volkmer and Heinemann, 2011).

We replicated the 200 experimental treatments (10 duckweed source sites *×* 10 microbe source sites *×* 2 zinc treatments) 8 times, and randomly arranged these 1600 experimental units across 67 24-well plates. We added 2.5 mL of sterile high- or low-zinc media, one mother-daughter frond cluster of surface-sterilized duckweed (dipped in 0.5% bleach for 30 s), and 80 *µ*L of microbial inocula to each well. We sealed plates with BreatheEasy membranes (Millipore-Sigma, Diversified Biotech, Dedham, MA, USA) to prevent microbial contamination and cross-contamination, while still allowing gas exchange. We placed plates in a growth chamber set to the same conditions as for the duckweed cultures (above), and let them grow for 14 days. All microbial or sterile manipulations were performed in a biological safety cabinet (ESCO Micro Pte. Ltd., Labculture*Q*_R_, Singapore). We autoclaved all media solutions prior to use.

We photographed plants with a custom-built camera rig (Nikon D3200 with AF-S DX NIKKOR 18-55mm f/3.5-5.6G VR lens, Minato, Tokyo, Japan, Yongnuo YN-300 light, Shenzhen, China) on day 0 and day 14, removing plate seals on day 14 for accurate color. We used ImageJ (Schneider et al., 2012) to measure plant growth and traits. We calculated the mm^2^ area of live duckweeds in each well, which is tightly correlated to the number of fronds (O’Brien et al., 2020). We averaged three measures across duckweed in each well: color intensity, aggregation (the ratio of frond area to frond perimeter, see O’Brien et al., 2019), and roundness of separate duckweed outlines (4 *× A/*(*π × d*^2^), where *d* is the major axis length, and *A* is the area). Aggregation becomes high when daughter fronds are retained on the mother (i.e., do not abscise), a process that varies across duckweed genotypes and in response to stress (Newton, 1977; Henke et al., 2011; O’Brien et al., 2019; O’Brien et al., 2020). Roundness may capture elements of both aggregation, as fronds that have not abscised have a less round outline, and frond expansion, which duckweed may also vary in response to environmental conditions (Paolacci et al., 2018). Color intensity quantifies brighter, or whiter pixels in images, and high values may indicate thinner fronds, capturing another axis of frond expansion.

After photographing on day 14, we froze and stored plates at −20^*°*^C. We then extracted and defrosted plates individually to measure optical density at 600 nm wavelength in 70 *µ*L from each well using a spectrophometer (BioTek Synergy HT with Gen5 1.10 software, Winooski, VT, USA). Optical density data was natural log-transformed, as microbial density is expected to be exponentially related to optical density (O’Brien et al., 2020). Microbial density is a measure of total microbial growth during the course of the experiment.

### Data analysis

#### Estimating plant genotype, microbiome, and environment effects

For all traits, we calculated the percent of variance explained by treatments as variance of the treatment means divided by total variance in the trait. We also calculated these percentages for duckweed frond area and microbial density. The variance explained by duckweed isogenic line can be interpreted as broad-sense heritability (e.g. Falconer, 1996, including additive, epistatic, and dominance genetic effects, as well as any transgenerational or epigenetic effects). The variance explained by the identity of the inoculated microbial community may be interpreted as the contribution of individual microbial strains, species, or their interactions (also in additive or non-additive ways) to duckweed fitness and phenotypes. The interactions of these terms with each other or with zinc treatment show how heritability in duckweeds or duckweed microbiomes depends on microbial or environmental context: e.g. *G*_*host*_ *× G*_*microbe*_, *G*_*host*_ *× E, G*_*microbe*_ *× E*, and *G*_*host*_ *× G*_*microbe*_ *× E* effects.

We assessed significance in linear models using the MCMCglmm package in R (Hadfield, 2010; R Core Team, 2019). In these models, we accounted for strong spatial effects by identifying the best model of spatial covariates for each response variable (x and y growth chamber coordinates, row and column plate coordinates, including both linear and squared terms). We then tested whether duckweed, microbiome, and zinc environments as well as their interactive effects explained significant variation in duckweed frond area, microbial density, and duckweed traits by testing whether adding duckweed and microbiome origin, zinc treatment, or their interactions as random effects (duckweed, microbe, duckweed*×*microbe, duckweed*×*zinc, microbe*×*zinc, duckweed*×*microbe*×*zinc) improved the fit over spatial effects alone.

In all linear models, we evaluated the significance of fixed effects by whether the highest posterior density intervals (HPDIs, Bayesian equivalents of confidence intervals) overlapped 0 at pMCMC *<* 0.5 (Bayesian equivalent of p-values), and if not, whether removing the term worsened model fit (evaluated by deviance information criterion, DIC, Spiegelhalter et al., 2002). Marginally significant terms (pMCMC *<*0.1, but *>*0.5) were retained in models as they generally improved fit, but were not considered ‘significant.’ Random effects cannot be estimated to be below 0, and so instead of using HPDIs, we assessed their significance solely by whether they improved DIC when included. For models of fixed effects, we removed non-significant terms one at a time and re-fit iteratively. For models of random effects, we iteratively added terms. All variables were fit with a Gaussian distribution. We used 11,000 MCMC iterations (increasing to 101,000 for selected best models), 1,000 burn-in, and thinned by 100.

#### Testing benefits of ecological or evolutionary history

We hypothesized that shared history of duckweed and microbes with each other or zinc should have predictable contributions to *G*_*host*_ *× G*_*microbe*_, *G*_*host*_ *× E, G*_*microbe*_ *× E*, and *G*_*host*_ *× G*_*microbe*_ *× E* effects. Local adaptation between duckweed and microbes and environmentally dependent local adaptation between duckweed and microbes would be sources of *G*_*host*_ *× G*_*microbe*_ and *G*_*microbe*_ *× E*, and *G*_*host*_ *× G*_*microbe*_ *× E* effects, respectively. Adaptation of duckweed or microbes from urban sites to zinc would be a source of *G*_*host*_ *× E* or *G*_*microbe*_ *× E* effects.

We first tested whether there was any evidence of local adaptation between duckweed and microbiomes (sympatric effects), and whether this was altered by zinc. We included spatial covariates as above. We also included random effects for duckweed line and microbe communities because genotypes that are always more fit or environments (microbes) that are always better can bias tests of local adaptation (Blanquart et al., 2013). Random effects are assumed to be drawn from normal distributions, *N*_*D*_(0) and *N*_*M*_ (0), respectively. Specifically, we fit the model *Y ∼ α*+*Sympatry*+*Zinc*+*Sympatry×Zinc*+*N*_*D*_(0)+*N*_*M*_ (0)+*f* (*Space*)+ *ϵ*, where *Y* is either duckweed frond area or microbial density (log of optical density at 600 nm) and *f* (*Space*) is the best fit spatial covariate effects described above.

Because microbiomes vary in composition, non-local microbiomes may differ more or less strongly from local microbiomes. Likewise, cultured microbiomes from the local site may vary in how accurately they represent the local field microbiome. Therefore, we also tested whether duckweed fitness (frond area) or microbial growth (microbial density, log of optical density at 600 nm) increased when inoculated microbiomes were more similar in composition to microbiomes observed on duckweed genotypes in the field (pairwise Unifrac distance, *PairwiseD*). Specifically, we fit the model *Y ∼ α* + *PairwiseD* + *Zinc* + *PairwiseD × Zinc* + *f* (*Space*) + *ϵ*, where Y is either duckweed frond area or microbial density.

Since particular taxa may sometimes contribute disproportionately to effects on hosts (Finkel et al., 2020), we sought to generate hypotheses about which bacterial groups might be stronger drivers of effects on host traits and fitness. We therefore conducted an exploratory analysis of associations between duckweed fitness or microbial growth and taxonomic components of the microbiome. For each phylogenetic balance calculated from the sequence data (relative abundance among subtending sister clades for nodes with sequences in *>*=8 inocula communities, see above), we recorded DIC of models for duckweed frond area and microbial density: *Y ∼ α* + *balance* + *Zinc* + *balance × Zinc* + *f* (*Space*) + *ϵ*.

Next, we tested whether microbes or duckweed sourced from more urban sites were better able to grow in high zinc, as we would expect if plants, microbes, or both were locally adapted to this urban aquatic stressor. We used the respective best-fit spatial effects coviariates model, and added effects of zinc treatment, the estimated percent of permeable surfaces (*PS*) within a 0.5 km radius (estimated with Google Earth imagery and ImageJ), and the interaction (*Y ∼ α* + *PS* + *Zinc* + *PS*: *Zinc* + *f* (*Space*) + *ϵ*, where *Y* is either response variable). Relatedly, we tested whether there was altered selective pressure on measured duckweed phenotypes in our different zinc contamination contexts by fitting models between our measure of duckweed fitness (frond area) and each trait, as well as between frond area and microbial density. We included linear and squared terms for the trait or microbial density, and the interactions of each with zinc environment: *Y ∼ α* + *Trait* + *Trait*^2^ + *Zinc* + *Trait × Zinc* + *Trait*^2^ *× Zinc* + *ϵ*.

### Testing whether more dissimilar microbiomes increase *G*_*host*_ *× G*_*microbe*_

We hypothesized that the effects of inoculated microbiomes might depend on how similar microbiomes were, specifically: that we are more likely to observe host-dependent effects of microbiomes when comparing more dissimilar microbiomes. If host-dependent effects of microbiomes do not exist, then the correlation of trait values expressed by duckweed genotypes in one microbiome to values expressed by duckweed genotypes in another microbiome should be *≈* 1. Conversely, if host-dependent effects of microbiomes are strong, the correlation between trait values expressed by duckweed genotypes in one microbiome and values expressed by duckweed genotypes in another microbiome will be substantially reduced. Correlations could theoretically be as low as −1, which would indicate that the relative trait values of the duckweed genotypes were exactly reversed across the two compared microbiomes.

We calculated the correlation across duckweed genotype means between microbiomes for all pairwise combinations, and we call this correlation metric ‘trait similarity’. We measured microbiome similarity as the pairwise Unifrac distance between cultured microbiomes. We then tested whether similar microbiomes produce more similar trait effects with a linear model (*TraitSim ∼ α* + *PairwiseD* + *ϵ*).

## Results

### Cultured microbiomes are less diverse than field microbiomes but share taxa

Microbial communities from the field are substantially more diverse than inocula: amplicon sequence variants (ASVs) totalled 3,773 across field samples versus 198 across cultured communities. Culturing communities also biased the taxonomic composition of experimental microbiomes. Some families that were common in sequence data from field-collected duckweed, such as *Rhodobacteraceae* and *Pirellulaceae* (1-28% and 1-23% of field samples), were very rare in cultured communities (detected in 1 community at less that 0.05%). Cultured communities were instead dominated by *Aeromonadaceae* and *Pseudomonadaceae* (Figure 1a), which were sequenced at low abundance in field communities (7 and 9 field samples, at a max of 0.5 and 0.9%, respectively). ASVs that overlapped between a cultured and the matching direct-sequenced field community rarely comprised a major proportion of cultured communities (Figure 1b, solid blue bars, 0-19%).

**Figure 1:**
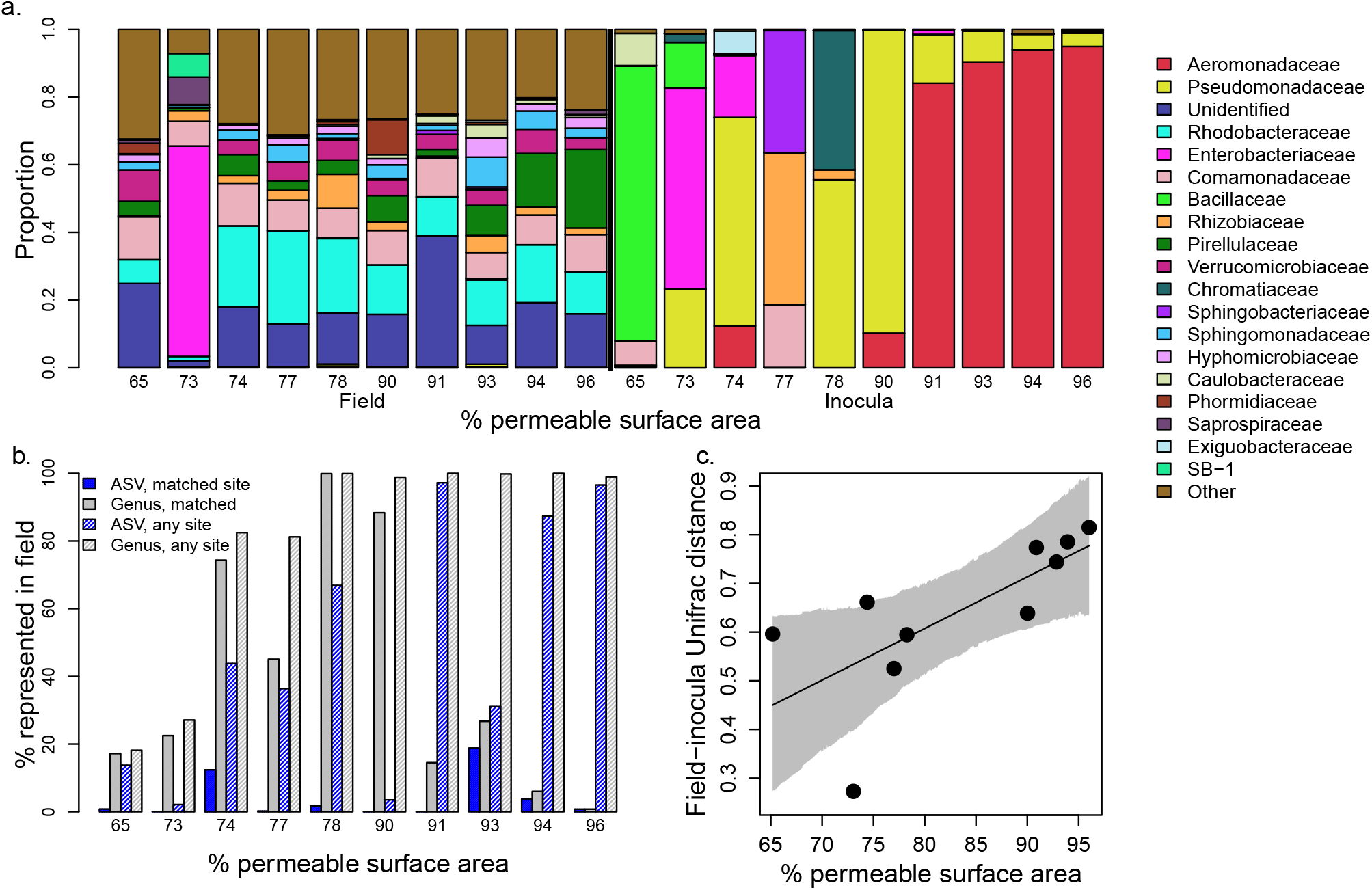
a) Relative abundance of 16s rRNA gene sequences identified to family. Bars are ordered by field (left 10 bars) or inoculum (right 10 bars) collection site and by percent permeable surface area within a 0.5 km radius (increasing from left to right). ‘Unidentified’ refers to the sum of all groups of sequences that could not be identified to family level. ‘Other’ refers to the sum of all groups that were individually *<* 5% of the total sequences. b) Proportion of identical sequences (blue) or proportion of sequences belonging to a genus (gray) that was also captured in a field sample – either any field sample (hatched) or the matched site (solid). c) Pairwise Unifrac distance (higher indicates communities are less phylogenetically similar, weighted by taxa abundance) between field and paired inoculum plotted against the the percent permeable surface area in a 0.5 km radius. Line and shaded region indicate the prediction and 95% HPDI for a linear model of field-inoculum Unifrac distance and percent permeable surface area.

Nonetheless, microbial communities isolated from duckweed from the field had some similarities to field samples. A sizable proportion of the sequences from cultured inocula communities belong to ASVs or identified genera captured in sequences from in at least one field site, though this is true mostly in microbiomes cultured from rural sites (Figure 1b, hatched blue and grey bars, 2-97% and 18-100%). Sequences from cultured inocula also often belonged to named genera that were captured in the directly sequenced field microbiome from the same site (Figure 1b, solid grey bars, 1-100%), though this was highest in our sites of intermediate urbanness. In contrast to ASV and genera relative abundance, Unifrac distances (accounting for both relative abundance and phylogenetic similarity) showed that field and matched cultured communities from urban sites were more similar overall than matched communities from rural sites (Figure 1c, p*<*0.05, see also S1), presumably due to higher abundance of families that were rare in field samples in these rural sites (*Aeromonadaceae, Pseudomonadaceae*).

### Variation in fitness, traits caused by treatments

Duckweed line (*G*_*host*_) was an important source of variance in trait values and was retained in all models as either a main random effect (explaining 24%, 22% and 7.5% of the variation in duckweed frond area, microbial density, and roundness, respectively), or as an interactive random effect (together with with microbial treatment, *G*_*host*_ *× G*_*microbe*_, explaining 27% and 24% of variation in color intensity and aggregation, respectively, Figure 2). For all response variables except frond roundness, microbiome treatment (*G*_*microbe*_) was identified as an important source of variation, either as an interactive random effect with duckweed line (above) or as an interactive random effect with zinc (*G*_*microbe*_ *× E*, explaining 1.8%, 1.5%, and 6.1% of duckweed frond area, microbial density, and aggregation). Zinc treatment (*E*) was not important alone, as it was not identified as an important main random effect, and it explained only a small amount of total variance (0-0.5%, Figure 2). We therefore observed important *G*_*host*_, *G*_*microbe*_, *G*_*microbe*_ *× E*, and *G*_*host*_ *× G*_*microbe*_ effects on fitness or trait expression in duckweed, indicating contributions of duckweed and microbes to broad-sense trait heritability, shifts in host contributions to heritability across microbiome contexts, and shifts in microbiome contributions to heritability across abiotic contexts.

**Figure 2:**
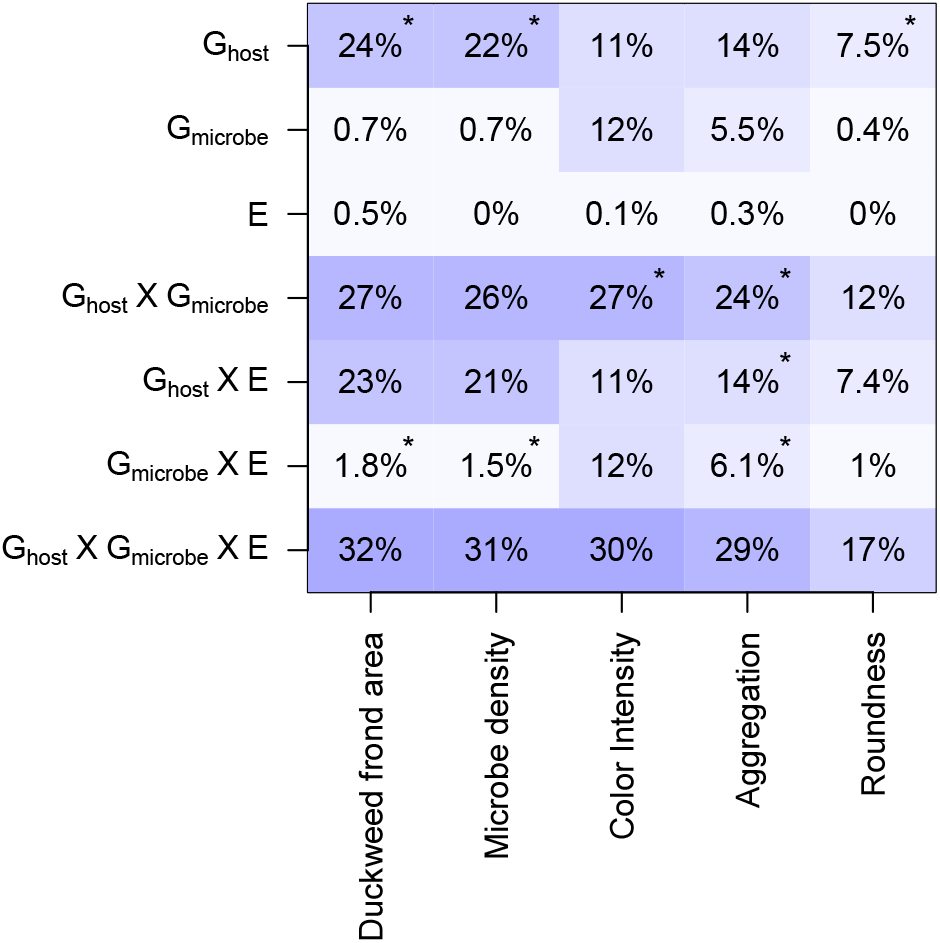
Percent of variance explained by treatments for duckweed fitness (frond area), each trait, and microbial density (log of optical density at 600 nm, a measure of total microbial growth). Asterisks (*) indicate significant random effects in linear models for each trait that also account for spatial effects. Background shading highlights larger percentages.

### Shared history contributes to *G*_*host*_ × *G*_*microbe*_ and *G*_*microbe*_ *× E* effects on fitness

Zinc negatively affected duckweed fitness, but had no impact on microbial growth (Figure 3). On average, duckweed frond area decreased by about 5% in the high-compared to low-zinc (from an average of 5.96 to 5.67 mm^2^ *±* a standard error of 0.10 for both). However, the effects of zinc on both duckweed frond area and microbial density were conditional on other contexts.

**Figure 3:**
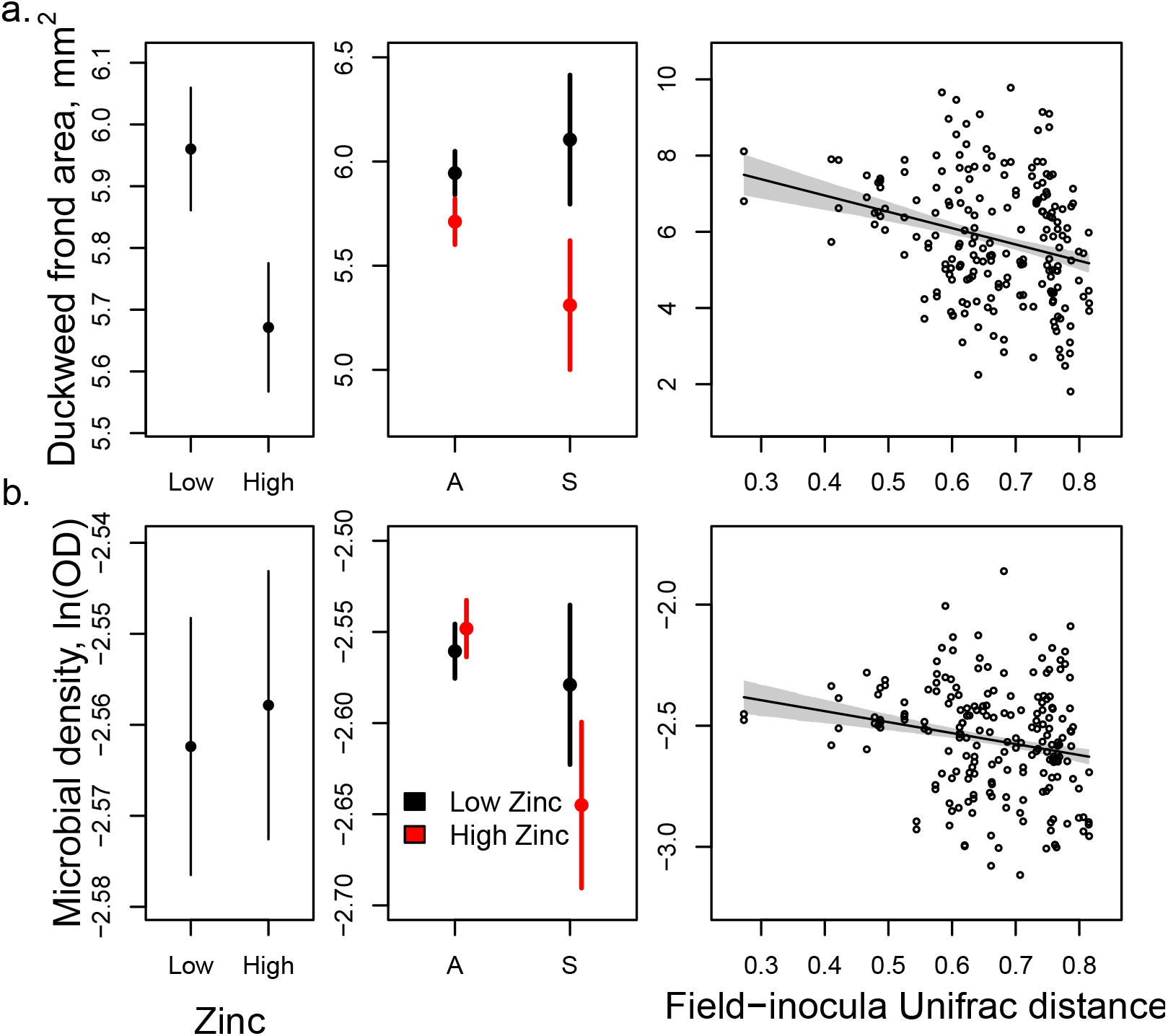
Left panels: Effect of zinc on duckweed fitness (frond area, top, significant at pMCMC *<* 0.05) and microbial density (log of optical density at 600 nm, a measure of total microbial growth, bottom, not significant). Means are shown in points and*±*one standard error of the mean as bars. Middle panels: Means and standard errors for duckweed and microbial growth in high (red) and low (black) zinc treatments for sympatric (‘S’) or allopatric (‘A’) combinations of duckweeds and microbes. Right panels: Relationship between duckweed frond area or microbial density and the pairwise Unifrac distance between the field microbiome of the duckweed host and the experimentally inoculated microbiome. Significance and model predictions in Tables S4 and S5.

Microbiomes cultured from the same sites as duckweed hosts, or microbiomes that were more similar to ‘home’ microbiomes, were not equivalent to other microbiomes in their effects on duckweed fitness. Local communities non-significantly increased duckweed frond area at low zinc conditions, but marginally decreased frond area at high zinc conditions (by about 8%, at 90% HPDI, Figure 3, see also Table S4). Similarly, the effect of inoculating duckweed with local versus non-local microbial communities affected microbial cell density only in high-zinc treatments, where it negatively impacted microbial growth (p *<* 0.05, Figure 3) with an average decrease in optical density from 0.078 to 0.072, corresponding to an expected reduction in cell density of about 3,200 cells/*µ*L (see also Table S5). On the other hand, cultured microbiomes that were more similar to field communities isolated from the duckweed host significantly increased the frond area of duckweed and microbial density (both pMCMC *<* 0.05). There was about a 30% increase for duckweed frond area from least to most similar microbiomes, and a shift in optical density of from 0.072 to 0.092, corresponding to an expected cell density increase of about 10,000 cells/*µ*L. There was no significant interaction between community similarity and zinc treatment.

Interactive random effects between microbiome and zinc treatments were not fully explained by sympatric versus allopatric comparisons, nor by similarity of communities to field microbiomes. Therefore, we considered whether particular subsets of taxa in cultured microbiomes are associated with duckweed fitness or microbial growth. Duckweed frond area was most strongly correlated to the relative abundance of sister clades within the *Pseudomonadaceae* found in inocula - but only in low zinc environments. In contrast, microbial growth was most correlated to the relative abundance of a clade including the *Rhizobiaceae* to a clade including the *Caulobacteriaceae*, but only in high zinc treatments (Figure S4).

### Zinc history does not alter zinc tolerance

Despite the negative effect of zinc on duckweed fitness, we saw little to no evidence of adaptation to high zinc conditions in duckweed collected from more urban sites. Effects of zinc on duckweed fitness and microbial growth did not depend on the % permeable surface area at the site from which they were collected. Duckweed from urban sites (low % permeable surface area) grew to much larger total frond area at high zinc, but also at low zinc (pMCMC *<* 0.05, Figure S3). Microbes from urban sites and microbes inoculated onto duckweed from rural sites reached higher cell density (pMCMC *<* 0.1 and 0.05, respectively, Figure S3), regardless of zinc treatments.

We also saw little evidence for differential selection on measured phenotypes in different zinc contamination contexts (Figure 4). Traits were variable across duckweed lines (Figure 2, see above), and were related to our measure of duckweed fitness, frond area. However, trait-fitness relationships were similar in high and low zinc treatments. For all duckweed traits and microbial density, the relationship to duckweed fitness was significantly curvilinear (negative parabolic terms, Table S6), suggesting that stabilizing selection may act on these traits. For color intensity, and aggregation, the optimal value was at the high end of the distribution, but for roundness and microbial density, the optimal value was mid-range. However, only the relationship between microbial density and duckweed changed significantly across zinc levels, and even this change represented only a slight decrease in the optimal amount of microbial density, with the distributions of fitnesses across microbial density largely overlapping (Figure 4, Table S6).

**Figure 4:**
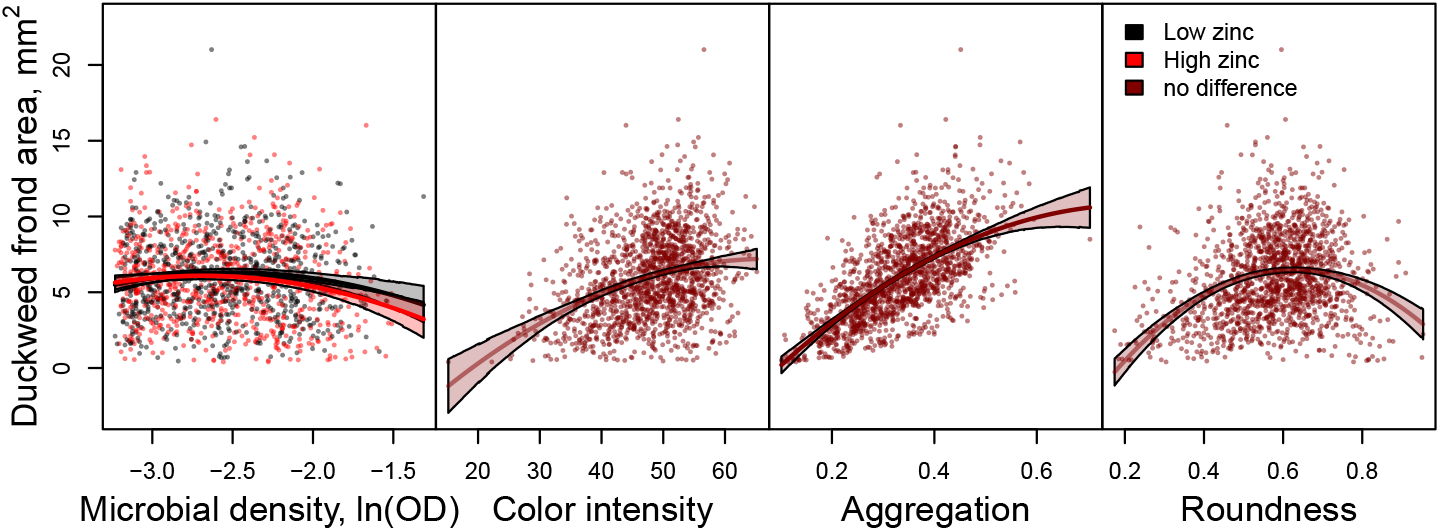
Relationships between duckweed fitness (frond area) and trait values or microbial growth. Each point is a single well of the experiment. Lines indicate predictions for the mean and shaded regions indicate 95% HPDI from models (see Table S6). Lines, shaded regions, and points are colored by zinc treatment (red for high zinc, black for low zinc) only for microbial density (log of optical density at 600 nm, a measure of total microbial growth). Other measures did not have significantly different trait-fitness relationships across zinc treatments, and lines, shaded regions, and points are all dark red.

### More dissimilar microbiomes drive *G*_*host*_ *× G*_*microbe*_ effects

The effect of duckweed line on fitness, trait, and microbial growth values remained similar only when comparing duckweeds grown in similar microbial communities. When the correlation of duckweed line means in one microbial community to line means another is high, this indicates that the relative effects of these microbial communities on duckweed were similar across all hosts genotypes, e.g., that *G*_*host*_ *× G*_*microbe*_ effects are weak for that pair of communities. Conversely, when the correlation of duckweed line means across two microbial communities is low, this suggests that the relative effects of these microbial communities on duckweed were not similar across duckweed hosts, e.g., that *G*_*host*_ *×G*_*microbe*_ effects are strong for that pair of communities. The correlation of duckweed line means between pairs of microbial communities (here, trait similarity) was significantly negatively related to the Unifrac distance between those microbial communities (Figure 5, for duckweed frond area, microbial density, aggregation, and color intensity pMCMC *<* 0.05, but not roundness). Trait similarity ranged from 0.83-0.97 for the most similar microbiomes, but from 0.36-0.72 for the least similar microbiomes (across fitness, microbial density, and traits, excluding roundness). This indicates that when two microbiomes are dissimilar, *G*_*host*_ *×G*_*microbe*_ effects are stronger. The biggest shift in trait similarity across microbiome similarity was for aggregation, which ranged from 0.92 (95% HPDI 0.79-1) for the most similar microbial communities to 0.36 (95% HPDI 0.26-0.46) for the least similar.

**Figure 5:**
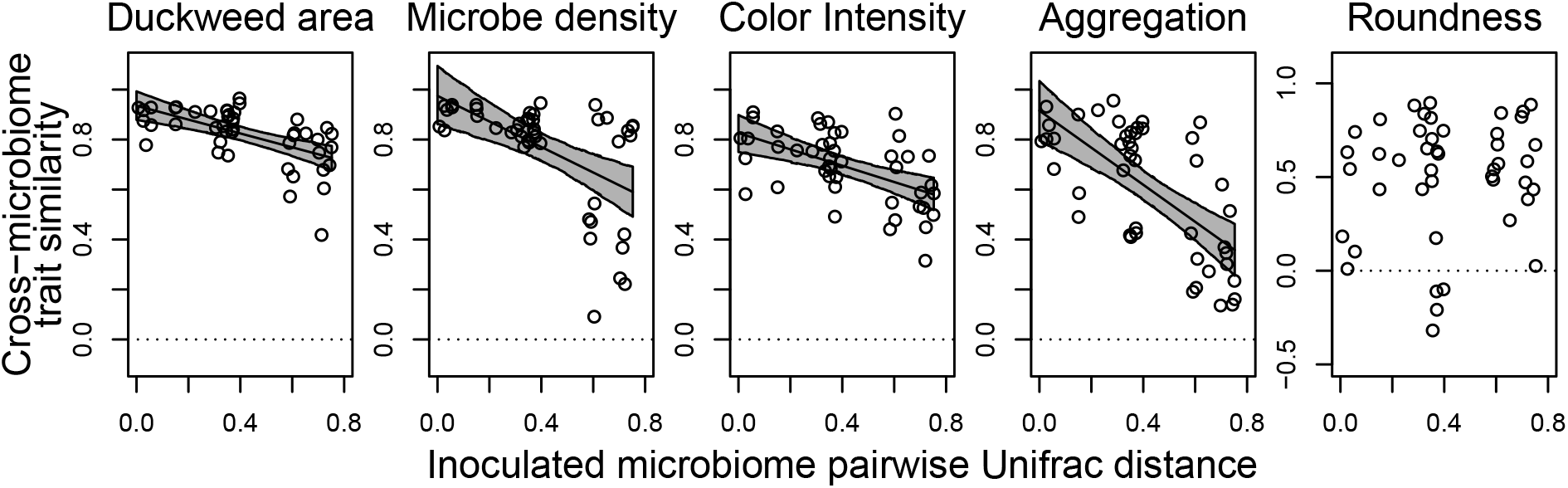
Correlations between pairwise similarity in trait or fitness genetic values estimated for the duckweed genotypes across each pair of microbial sources (y-axis) and microbial community distance of those same sources (x-axis). Plots from left to right represent different fitness and trait measures: duckweed frond area (duckweed fitness), microbial density (log of optical density at 600 nm, a measure of total microbial growth), duckweed frond color intensity, duckweed area:perimeter ratio (aggregation), and duckweed frond roundness. Lines and shaded regions represent predicted mean and 95% highest posterior density intervals for significant relationships (all but frond roundness, where this relationship is n.s., see Table S3). See also Figure S2, where each point here is expanded to a full panel.

## Discussion

A focus of recent host-microbiome research has been to determine whether microbiome community composition is heritable (Peiffer et al., 2013; Grieneisen et al., 2021), and whether this heritable variation in hosts responds to selection (Henry et al., 2021). Yet, if selection on hosts directly impacts the relative fitness of different species or strains, responses of microbiomes could contribute to trait change in hosts without any shift in host genotypes: via either ecological shifts in species composition (Lau and Lennon, 2012), or evolutionary changes within individual microbes (Batstone et al., 2020; Tso et al., 2018, see also Rebolleda-Gómez et al., 2019). Whether responses to selection on host traits include shifts in microbiome contributions is expected to hinge only on the amount of host phenotypic variation encoded by microbiomes (O’Brien et al., 2021), and our ability to engineer microbiome variation for desired effects on host traits will depend on our ability to predict microbiome effects on host phenotypes. Here, we quantified variation in duckweed fitness and phenotypes that is attributable to the manipulable fraction of their microbiome. We found that microbiomes contribute to traits, but primarily via host- or environment-dependent mechanisms.

### Host genotype underlies most variation, but effects are only predictable in similar microbiomes

Duckweed line was the largest source of phenotypic variation, though it sometimes depended on which microbiome was inoculated onto plants. Other studies have also found that most host trait variation is explained by host genotype, or host-specific impacts of microbes (Wagner et al., 2014; O’Brien et al., 2019), though few studies have attempted to evaluate microbial effects across both multiple host genotypes and multiple environments (but see Fitzpatrick et al., 2019).

In contrast, microbiome contributions to host traits and fitness were always contingent on host genotype or environment (i.e., zinc). The context-dependency of microbe effects could result from several processes. Host genotypes may affect the relative growth and success of different taxonomic groups within inoculated communities, resulting in different assembled microbiomes from the same source microbial communities (Lebeis et al., 2015; Jackrel et al., 2021; Wippel et al., 2021; Grieneisen et al., 2021). Differential assembly across host genotypes could result from top-down host selection of different microbes (Lebeis et al., 2015), or from adaptation of microbes to local hosts in colonization ability (Li et al., 2021). Abiotic environments can also alter microbiome assembly (Wang et al., 2021), as microbiome members often have differential tolerance to stressors, including zinc (Davis et al., 2004; Jain et al., 2020), which could account for zinc-dependent effects of microbiomes on duckweed fitness and traits.

Phenotypes resulting from host and microbiome variation were somewhat predictable from microbiome composition: more similar microbiomes resulted in conserved effects of host genotypes on traits, but less similar microbiomes resulted in unpredictable phenotypes across duckweed genotypes (Figure 5). This pattern suggests that for sufficiently similar communities, host-by-microbiome interactive effects (*G*_*host*_ *× G*_*microbe*_) effects may be minimal. When engineering small shifts to synthetic microbiome communities, we may be able to model microbiome effects as largely additive effects on host traits without contingent effects (e.g. Falconer, 1996; Finkel et al., 2020), which could simplify efforts to construct beneficial microbiomes. Reciprocally, this result also suggests that even host genotypes that perform poorly in one microbial condition may perform well in a very different microbial condition.

### Any benefits of shared history between duckweed and microbiomes disrupted by zinc

In other systems, plants often perform better with microbes from their local site, suggesting local adaptation between plants and microbiomes is common (Rúa et al., 2016). Furthermore, experimental evolution has documented adaptation of microbes to increase both host and microbe fitness in local combinations (Batstone et al., 2020). Here, the only significant difference between ‘home’ and ‘away’ microbes was at high-zinc, and revealed local maladaptation; in high-zinc conditions, combining duckweed and microbes from the same site reduced the fitness of duckweed and the growth of microbes.

Several mechanisms could explain this result. First, only mutually stressful environments are expected to strengthen mutualisms (Bronstein, 1994; O’Brien et al., 2018), and zinc is a stressor for duckweed, but not microbes (O’Brien et al., 2020, and Figure 3). Novel environmental conditions that are not mutually stressful, such as zinc, may be more likely to disrupt positive interactions. For example, nutrient pollution with positive effects on the fitness of one partner but not the other can disrupt or weaken the benefits of interactions (Shantz et al., 2016), leading to less mutualistic evolutionary trajectories (Klinger et al., 2016). Alternatively, cultured sympatric microbiomes may underestimate benefits received by duckweed from local microbiomes in the field. Furthermore, comparison between synthetic microbiomes cultured from the same or different sites may be insufficient to demonstrate local adaptation if culturing introduces composition shifts in microbiomes – e.g. because this could mean that ‘home’ microbiomes are not always the most similar microbiome to the field microbiome (Figures S1, 3).

### A closer ‘match’ to the natural microbiome matters for both hosts and microbes

We saw that microbiomes more similar to the community present on duckweed lines in the field consistently supported greater duckweed fitness and grew more themselves (Figure 3), and while there is overlap between cultured and field communities, there are major differences (Figure 1). If this is a general pattern, other studies of cultured ‘synthetic’ microbiomes may similarly not be testing the most beneficial microbiomes. It also suggests that studies evaluating single microbes outside of their community context, or using a culturing or storage step may underestimate whether local microbes are beneficial. Current estimates of the frequency of local adaptation in plant-microbe interactions (e.g., Rúa et al., 2016) could be underestimates.

The effects of microbial communities on hosts are not often compared to how well each community reflects the communities that are present on hosts in the field. We expect that inocula more similar to field microbiomes are more beneficial because the culture step may select for bacteria that grow well outside the host environment, whereas the most beneficial bacteria likely grow best in the host, and less well in culture. For example, Panke-Buisse et al. (2017) found that conditions in an intervening culture or storage phase drastically influenced microbiome effects on host traits, and Burghardt et al. (2018) found that microbial fitness in the host is only weakly correlated to microbial fitness in lab culture. This suggests that many beneficial effects of microbiomes are lost during sampling and culture, and that microbiome scientists should sample inocula more similar to field microbiomes. While it remains challenging, novel methods are helping to reduce bias in sampling and culturing (Thrash, 2021).

### Urban duckweed and microbes are not adapted to tolerate zinc

We found no evidence that urban duckweed or microbes are better adapted to higher zinc conditions: while both duckweed and microbes sourced from urban sites grew better in our experiments, this did not depend on zinc treatment (Figure S3). As we observed no effect of zinc on microbial growth (Figure 3), we would not necessarily expect any adaptive response, although other work on microbial zinc tolerance has found that prolonged exposure to high zinc can cause acquired zinc tolerance in microbial communities (Davis et al., 2004). For duckweed we had a stronger *a priori* expectation to observe adaptation to zinc: zinc is a known stressor for duckweed (O’Brien et al., 2020, see Figure 3), and adaptation to increased heavy metals including zinc has been documented in various plants (Antosiewicz, 1992; Schvartzman et al., 2018).

Several explanations for a lack of acquired zinc tolerance are possible. First, there may be no genetic variation for zinc tolerance in urban duckweeds. While relationships between most of our measured traits and duckweed fitness do not shift across zinc (Figure 4), duckweed lines did differ in the average effect of zinc (Figure S5). This suggests that there is variation in zinc tolerance in duckweed, but that it is unrelated to the traits we measured here. Instead, zinc exposure may not be uniformly higher in urban environments: urban sites range two orders of magnitude in zinc concentrations (Göbel et al., 2007), and some rural sites can receive high zinc runoff due to agricultural pesticides and fertilizers (Zhang et al., 2003). Another likely explanation is that urban environments include stressors beyond zinc. Urban environments have many concomitant differences from rural environments (Johnson and Munshi-South, 2017), and urban stormwater in particular is often a complex mix of synthetic organic compounds and other inputs novel to ecosystems (Masoner et al., 2019). The best strategy to persist may simply be to grow and reproduce quickly: like our duckweed, faster growth or shorter time to reproduction in genotypes from urban areas has been observed in other species (Brans and De Meester, 2018; Gorton et al., 2018; Santangelo et al., 2020, see Figure S3).

## Conclusions

While plant genotype harbours most variation for plant traits, microbes do alter traits, fitnesses, and their heritability, in host- and environment-dependent ways. The fact that *G*_*host*_ *×G*_*microbiome*_ effects are lower for similar microbiomes may simplify evolutionary models of these effects, and may increase predictability of synthetic microbiome effects. Actually releasing engineered microbiomes into the wild could have negative unintended consequences on environments and ecosystem services (Jack et al., 2021), and we will need to understand the potential ecological and evolutionary consequences of engineered microbiomes to guard against this possibility. Our results here suggest that not all synthetic microbiomes will be equally beneficial for hosts, and that synthetic microbiomes that are very different from what a host encounters in nature may reduce host fitness and lead to unpredictable host traits. Greater emphasis on culturing and testing synthetic microbiota that better match natural microbial communities is needed in microbiome science.

## Acknowledgements

The project was funded by the Gordon and Betty Moore Foundation through Grant GBMF9356 to MEF (https://doi.org/10.37807/GBMF9356) and an NSERC Discovery Grant to MEF. The authors would like to thank E. Lash for help collecting duckweeds, students and volunteers who have contributed to maintaining duckweed and microbe cultures in the lab, and members of the Frederickson lab for useful discussion.

---

**Table S1:** Collection locations for duckweeds and microbes in longitude (Lon) and latitude (degrees north, Lat N). Also provided are the calculated distance from the city center of Toronto, Ontario, Canada, and the percent permeable surface area in a 0.5 km radius of the point.

| Site name | Lon | Lat N | km to Toronto center | % Permeable surface area |
| --- | --- | --- | --- | --- |
| Arsandco Pond | -79.2672 | 43.7481 | 14.1 | 65.2 |
| Broken Bridge | -78.9782 | 44.1200 | 61.3 | 90.8 |
| Cambelleville | -79.9755 | 43.4934 | 51.0 | 90.0 |
| Cedarvale Pond | -79.4194 | 43.6897 | 5.0 | 73.0 |
| Humber Bay Park East | -79.4772 | 43.6203 | 8.4 | 77.0 |
| Koffler Scientific Reserve | -79.5178 | 44.0350 | 43.9 | 93.9 |
| Moccasin Trail | -79.3319 | 43.7333 | 9.8 | 78.3 |
| Rattray | -79.6046 | 43.5181 | 23.3 | 92.3 |
| University of Toronto Mississauga | -79.6632 | 43.5472 | 25.5 | 74.4 |
| West Duffins | -79.2011 | 43.9922 | 40.5 | 96.0 |

**Table S2:**
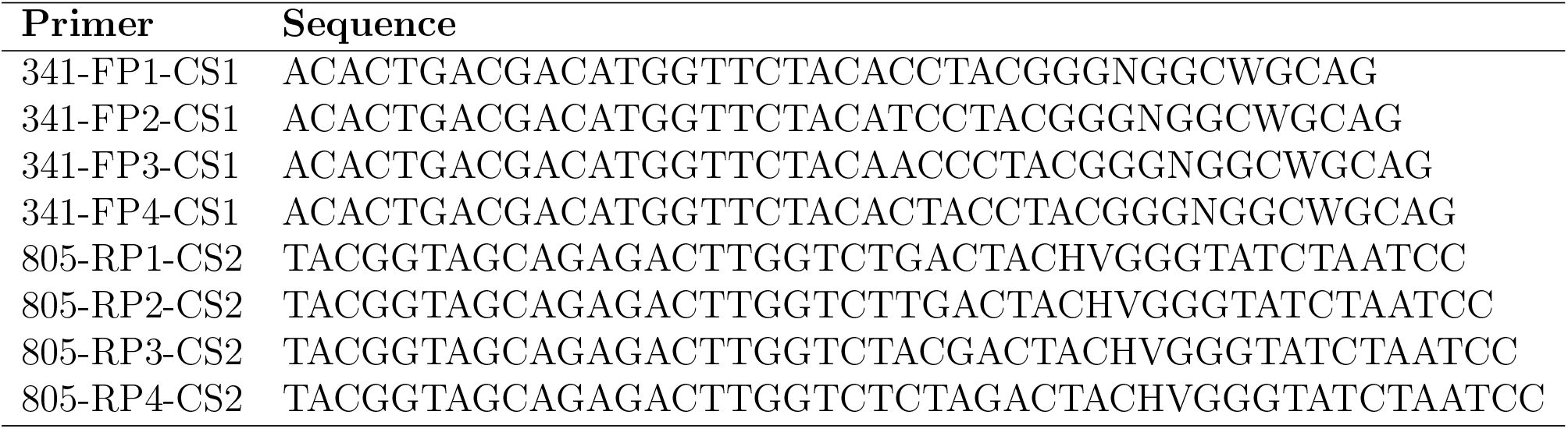
Forward and reverse primers used in amplification of the V3-V4 16S rRNA gene region.

**Table S3:** Fitted relationships between similarity of duckweed line means between each microbiome treatment and the Unifrac distance between those microbiomes. The response variable is the correlations of the average duckweed line trait values across each pair of microbiomes. Each row corresponds to one model, and the trait is listed in the first column. The density of microbial cells estimated by the log of optical density at 600 nm is our measure of microbial growth. ‘*’ indicates p *<* 0.05, ‘.’ indicates marginal significance, or p *<* 0.1.

| Trait/Fitness parameter | Intercept | Unifrac Distance |
| --- | --- | --- |
| **Duckweed frond area mm<sup>2</sup>** | 0.94* | -0.28* |
| **Microbial growth, ln(OD600)** | 0.97* | -0.51* |
| **Color intensity** | 0.83* | -0.33* |
| **Aggregation** | 0.92* | -0.74* |
| **Roundness** | 0.44* | 0.17 |

**Figure S1:**
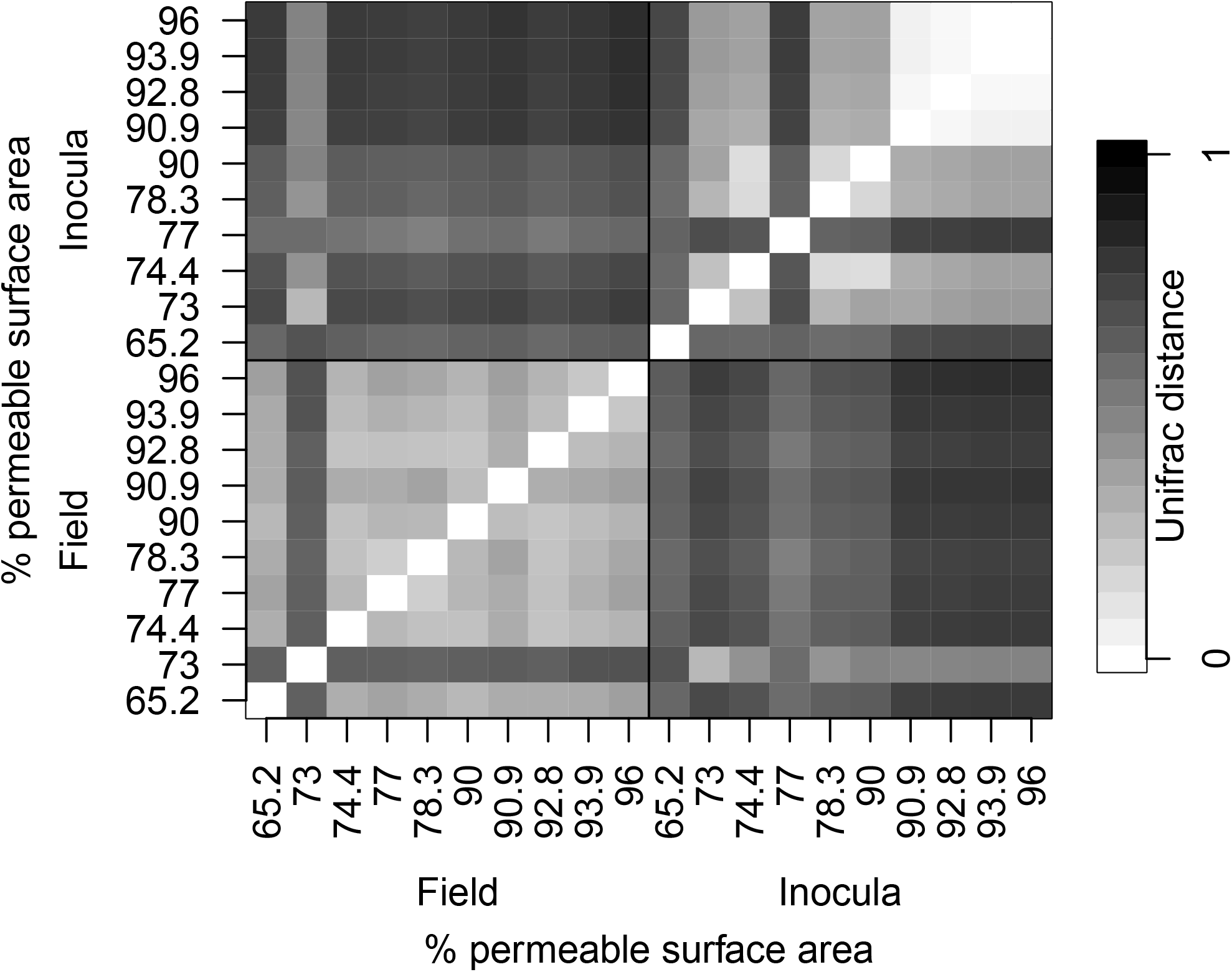
Pairwise weighted unifrac distance between all sequenced microbial communities. Sequenced communities from fresh duckweed and those from inocula are compared both within each category and to each other. Within categories, sites are ordered by increasing % permeable surface area. Darker squares indicate greater Unifrac distance and less similar communities, whereas whiter squares indicate more similar communities. The plot is symmetric around the diagonal.

**Figure S2:**
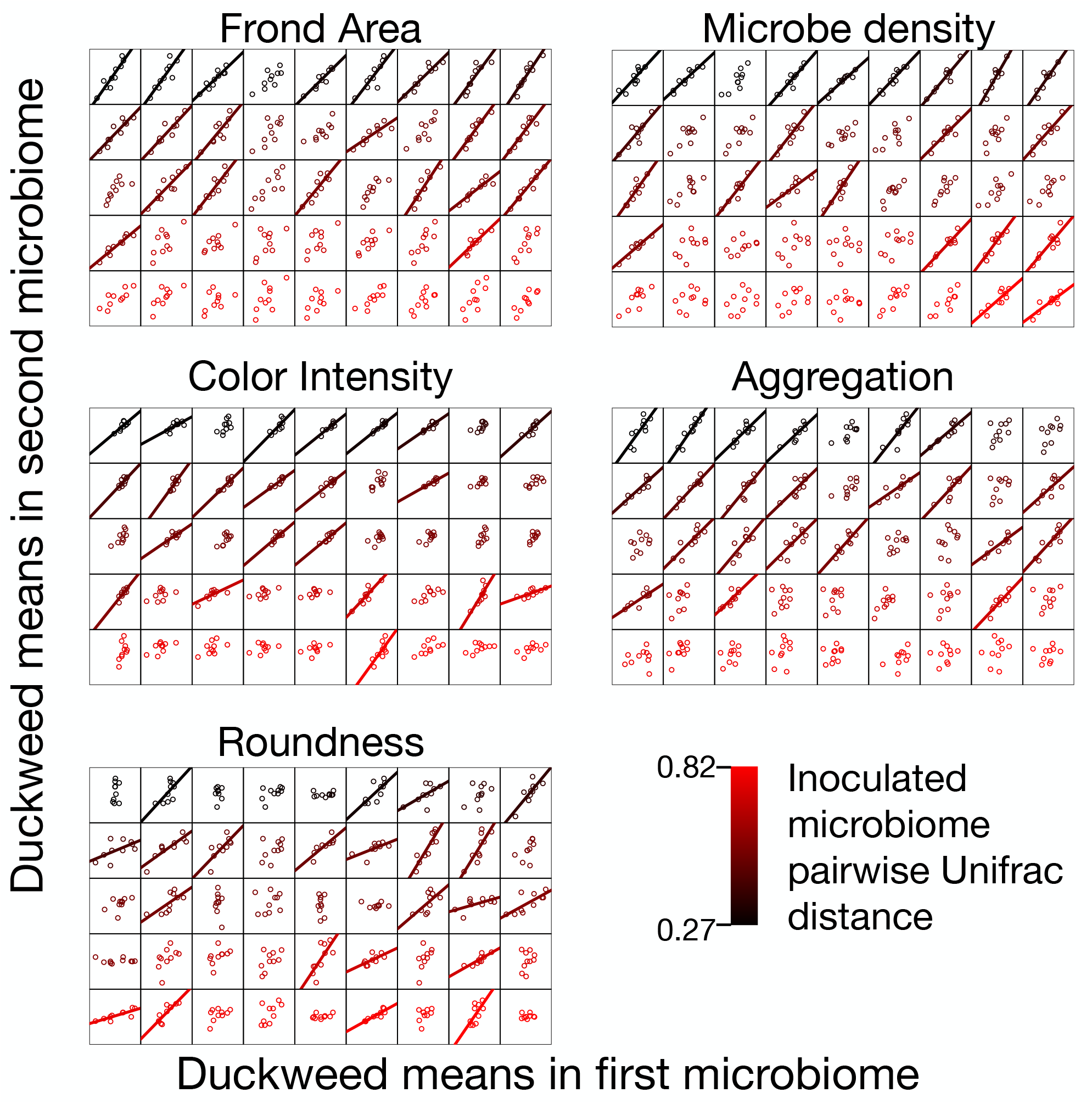
Each tiny panel is trait, fitness, or microbe density means of the 10 duckweed lines in one microbe plotted against their means in another. Each tiny panel represents one point in the corresponding panel of Figure 5. Plots within a trait are ordered by increasing Unifrac distance between the microbiomes (more distant = red). For each trait, trend lines are plotted for the top 50% correlations (22 of 45), between the trait means in the two different microbiome communities.

**Figure S3:**
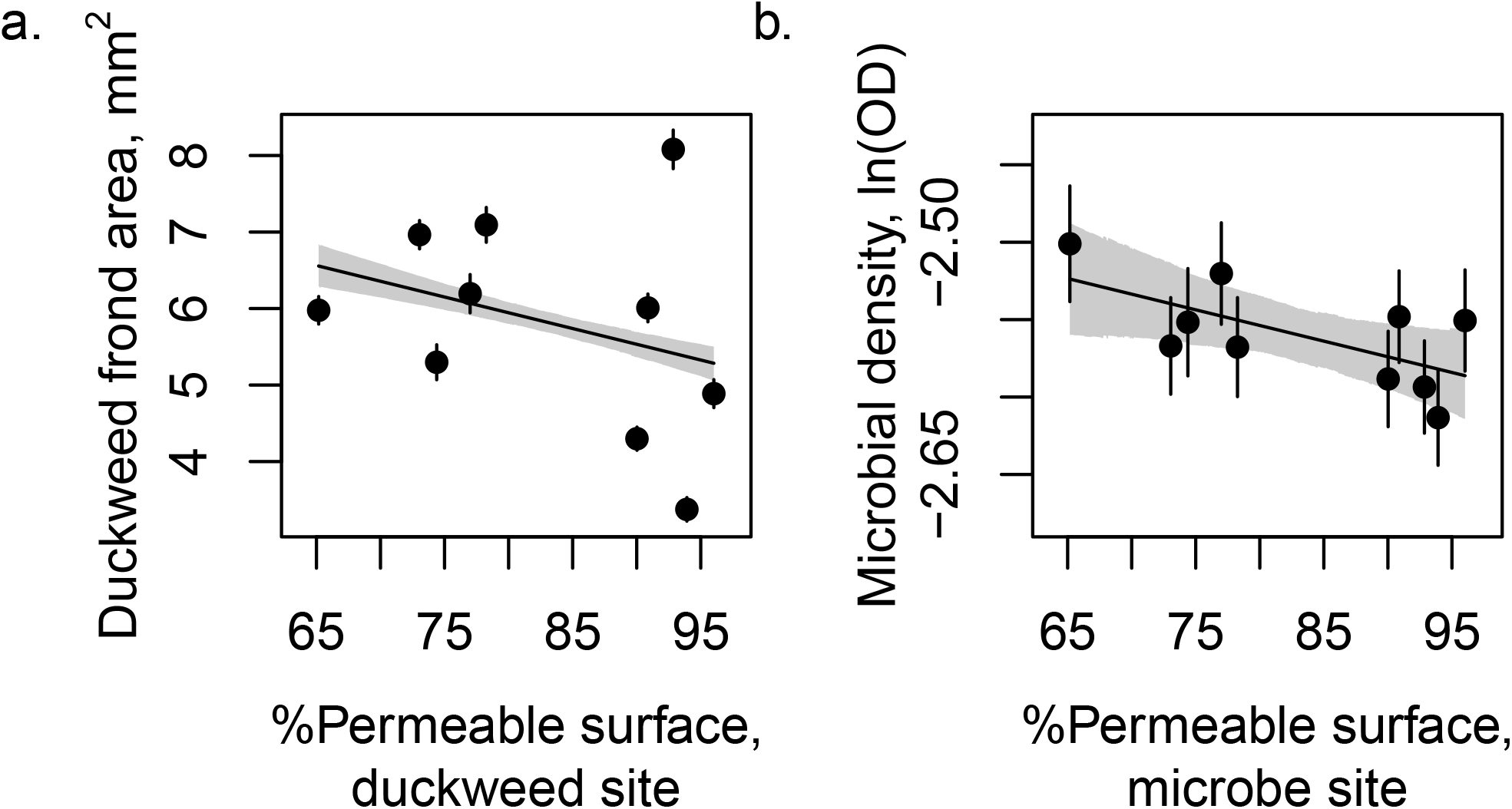
a) fitted relationship between our measure of duckweed final area and % of surface 0.5 km around the population origin that is not impervious (roofs, roads, sidewalks). Duckweed starting size has no relationship to the % of permeable surface around the source site. b) fitted relationship between our measure of microbial growth (log of the optical density at 600 nm) of surface 0.5 km around the population origin that is not impervious (roofs, roads, sidewalks). Means for each population by treatment average are shown as points with standard error bars, with predicted relationships and 95% HPDIs from the fitted model as lines and shaded regions, respectively.

**Figure S4:**
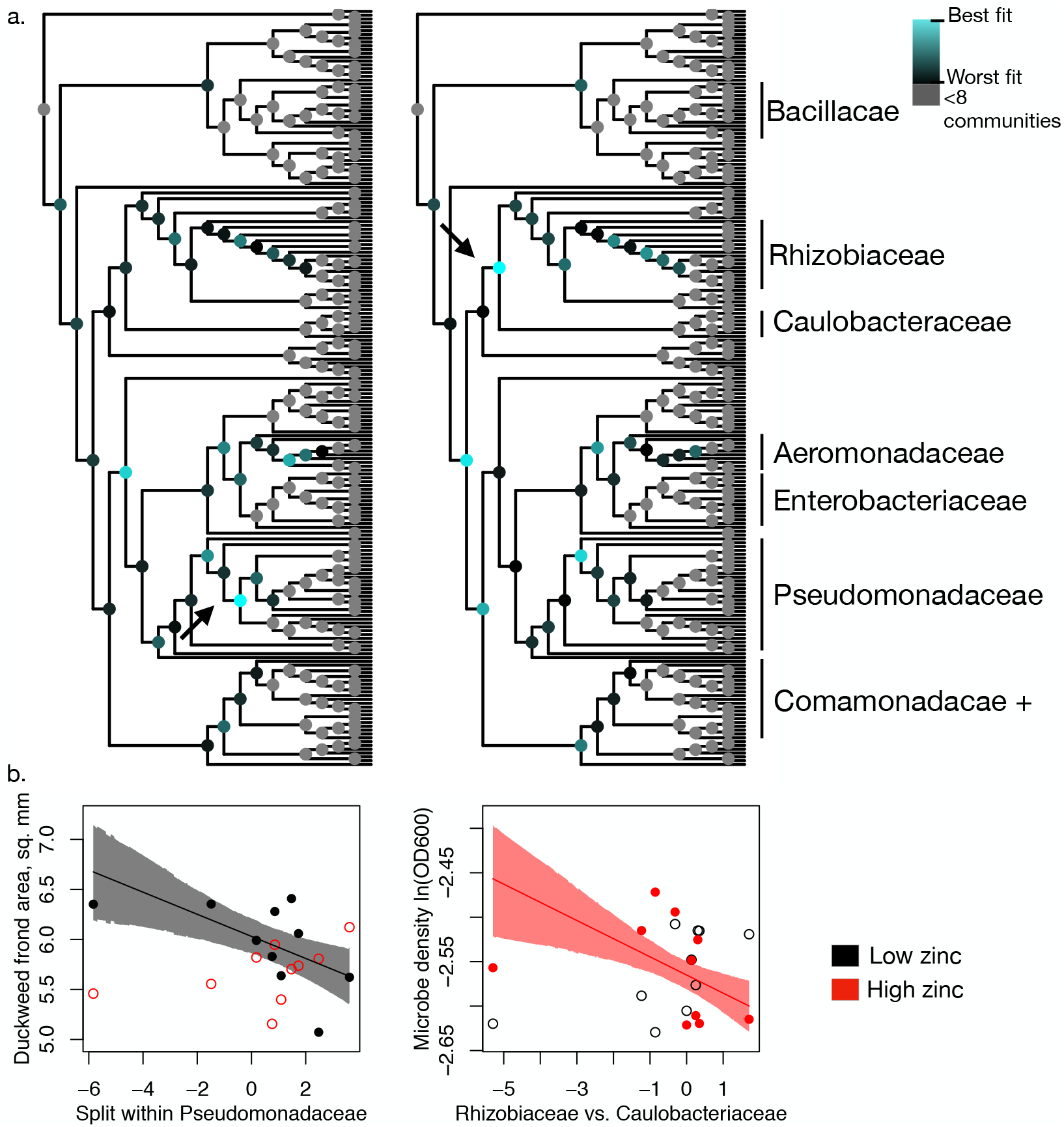
a) DIC for models between duckweed frond area (left) or microbial density (optical density at 600 nm, a measure of total microbial growth, right) and balances across a node (relative abundance ratio of summed daughter taxa on each branch) computed at each node where a subtending taxon was sequenced from at least 8 inocula communities. Fit between the balance and duckweed frond area or microbial growth is marked on nodes: lower DIC (better fit) is in blue, higher DIC in black, nodes where all subtending taxa are missing in more than 1 inoculum are in grey. Select families marked. While DIC for each node was computed for three model structures (depending on the inclusion of main and interactive effects with zinc treatments), only one is presented for each response variable (including all effects for duckweed frond area, and including only the interactive effect of the balance and zinc for microbe density) b) linear models for the most related balances for duckweed frond area and microbial density – correspond to nodes highlighted by arrows in trees in top plot. Note that this analysis is meant to be hypothesis generating and significance tests are not appropriate.

**Table S4:**
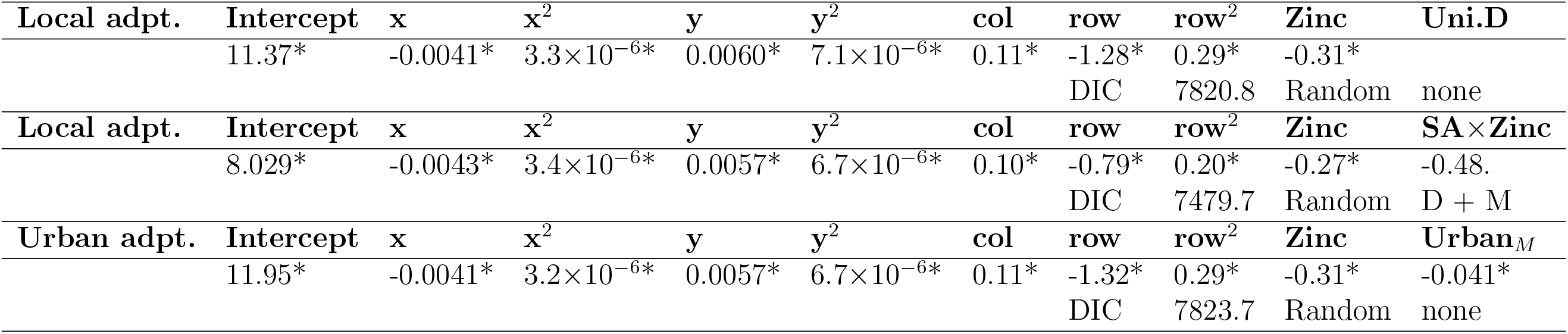
Hypothesis testing models fitted to duckweed frond area, including spatial effects. The first column of the table describes the *G*x*G* or *G*x*E* hypothesis tested by the model. ‘Uni.D’ refers to weighted pairwise Unifrac distance between the host duckweed’s field microbiome and inoculated microbiome. ‘SA’ refers to the fitted effect for sympatric or allopatric treatments, where sympatry is coded as 1 and allopatry as 0. ‘Urban_*D*_’ and ‘Urban_*M*_’ refer to the % permeable surface area at the collection site for the duckweed host and microbiome inocula, respectively. ‘col’ refers to the plate column for the well in the experiment, ‘row’ refers to the row in the plate. ‘x’ and ‘y’ are spatial coordinates of the experimental well in the growth chamber. Random effects ‘D’ and ‘M’ refer to duckweed line and microbial inocula, respectively. ‘*’ indicates p < 0.05.

|  |  |  |  |  |  |  |  |  |  |  |
| --- | --- | --- | --- | --- | --- | --- | --- | --- | --- | --- |
| **Local adpt.** | **Intercept** | **x** | **x<sup>2</sup>** | **y** | **y<sup>2</sup>** | **col** | **row** | **row<sup>2</sup>** | **Zinc** | **Uni.D** |
| | 11.37* | -0.0041* | $3.3 \times 10^{-6}$* | 0.0060* | $7.1 \times 10^{-6}$* | 0.11* | -1.28* | 0.29* | -0.31* | |
|  |  |  |  |  |  |  | DIC | 7820.8 | Random | none |
| **Local adpt.** | **Intercept** | **x** | **x<sup>2</sup>** | **y** | **y<sup>2</sup>** | **col** | **row** | **row<sup>2</sup>** | **Zinc** | **SA × Zinc** |
| | 8.029* | -0.0043* | $3.4 \times 10^{-6}$* | 0.0057* | $6.7 \times 10^{-6}$* | 0.10* | -0.79* | 0.20* | -0.27* | -0.48. |
|  |  |  |  |  |  |  | DIC | 7479.7 | Random | D + M |
| **Urban adpt.** | **Intercept** | **x** | **x<sup>2</sup>** | **y** | **y<sup>2</sup>** | **col** | **row** | **row<sup>2</sup>** | **Zinc** | **Urban<sub>M</sub>** |
| | 11.95* | -0.0041* | $3.2 \times 10^{-6}$* | 0.0057* | $6.7 \times 10^{-6}$* | 0.11* | -1.32* | 0.29* | -0.31* | -0.041* |
|  |  |  |  |  |  |  | DIC | 7823.7 | Random | none |

**Table S5:**
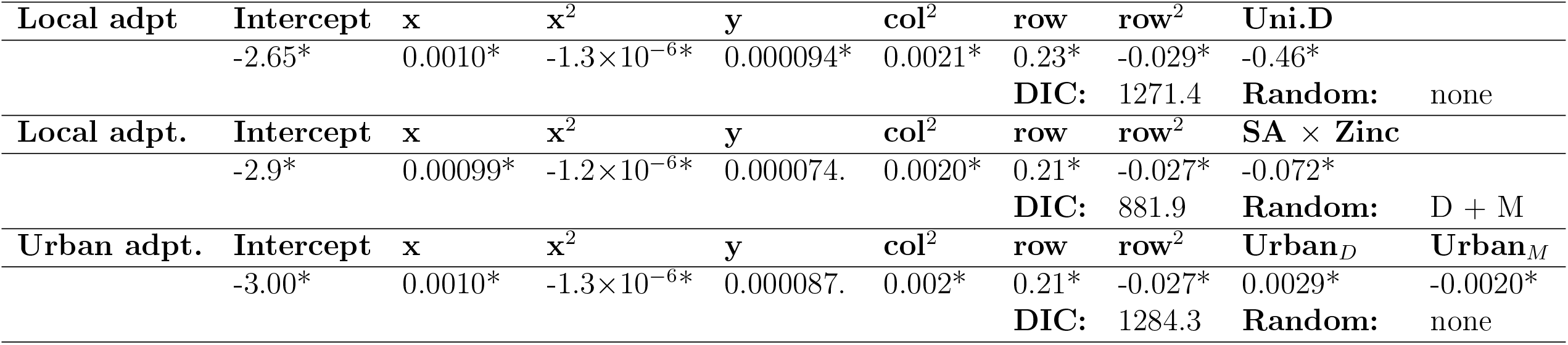
Hypothesis testing models fitted to microbial density (log of optical density at 600 nm, a measure of total microbial growth), including spatial effects. The first column of the table describes the *G*x*G* or *G*x*E* hypothesis tested by the model. ‘Uni.D’ refers to weighted pairwise Unifrac distance between the host duckweed’s field microbiome and inoculated microbiome. ‘SA’ refers to the fitted effect for sympatric or allopatric treatments, where sympatry is coded as 1 and allopatry as 0. ‘Urban_*D*_’ and ‘Urban_*M*_’ refer to the % permeable surface area at the collection site for the duckweed host and microbiome inocula, respectively. ‘col’ refers to the plate column for the well in the experiment, ‘row’ refers to the row in the plate. ‘x’ and ‘y’ are spatial coordinates of the experimental well in the growth chamber. Random effects ‘D’ and ‘M’ refer to duckweed line and microbial inocula, respectively. ‘*’ indicates p < 0.05, ‘.’ indicates marginal significance, or p < 0.1.

**Table S6:** Testing relationships between traits and duckweed fitness (frond area). Microbial growth refers to the log of media optical density at 600 nm at the final timepoint. Empty cells were parameters that were n.s. and removed from models. Each row is the best model using the trait given in the first column as the explanatory ‘Trait’ in the model. Duckweed frond area is the response for all models. ‘*’ indicates p *<* 0.05, ‘.’ indicates marginal significance, or p *<* 0.1.

| | Intercept | Trait | Trait<sup>2</sup> | Zinc | Zinc × Trait<sup>2</sup> |
| --- | --- | --- | --- | --- | --- |
| Microbial growth | -3.24 | -7.63* | -1.53* | -1.68 . | -0.56314 . |
| Aggregation | -3.10* | 34.86* | -21.88* |  |  |
| Color intensity | -6.83* | 0.42* | -0.0031* |  |  |
| Roundness | -6.48* | 41.44* | -33.13* |  |  |

**Figure S5:**
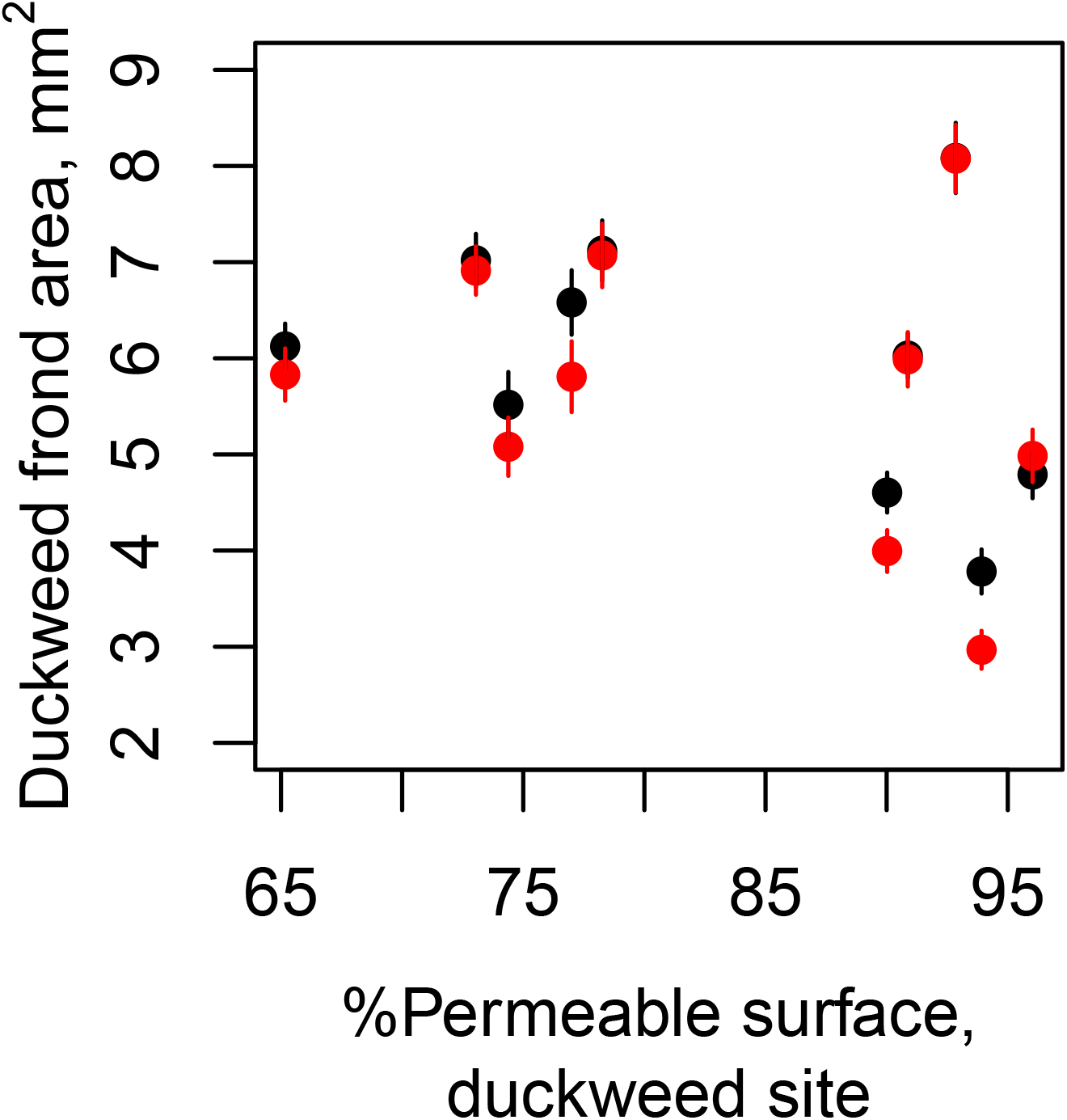
Average frond area of duckweed genotypes at high (red) and low (black) zinc. Points are means for each duckweed line and zinc treatment combination, and vertical bars are *±* one standard error of the mean. Some genotypes are more strongly negatively affected than others, implying genetic variation for zinc tolerance. Duckweed lines are plotted against the % permeable surface within a 0.5 km radius.

## References

Adler, L. S., R. E. Irwin, S. H. McArt, and R. L. Vannette. 2021. Floral traits affecting the transmission of beneficial and pathogenic pollinator-associated microbes. Current Opinion in Insect Science, 44:1–7.

Anderson, J. T., C.-R. Lee, C. A. Rushworth, R. I. Colautti, and T. Mitchell-Olds. 2013. Genetic trade-offs and conditional neutrality contribute to local adaptation. Molecular Ecology, 22:699–708.

Antosiewicz, D. M. 1992. Adaptation of plants to an environment polluted with heavy metals. Acta Societatis Botanicorum Poloniae, 61:281.

Batstone, R. T., A. M. O’Brien, T. L. Harrison, and M. E. Frederickson. 2020. Experimental evolution makes microbes more cooperative with their local host genotype. Science, 370:476–478.

Bertness, M. D. and R. Callaway. 1994. Positive interactions in communities. Trends in Ecology & Evolution, 9:191–193.

Blanquart, F., O. Kaltz, S. L. Nuismer, and S. Gandon. 2013. A practical guide to measuring local adaptation. Ecology Letters, 16:1195–1205.

Brans, K. I. and L. De Meester. 2018. City life on fast lanes: Urbanization induces an evolutionary shift towards a faster lifestyle in the water flea Daphnia. Functional Ecology, 32:2225–2240.

Bronstein, J. L. 1994. Conditional outcomes in mutualistic interactions. Trends in Ecology & Evolution, 9:214–217.

Buffington, S. A., G. V. Di Prisco, T. A. Auchtung, N. J. Ajami, J. F. Petrosino, and M. Costa-Mattioli. 2016. Microbial reconstitution reverses maternal diet-induced social and synaptic deficits in offspring. Cell, 165:1762–1775.

Burghardt, L. T., B. Epstein, J. Guhlin, M. S. Nelson, M. R. Taylor, N. D. Young, M. J. Sadowsky, and P. Tiffin. 2018. Select and resequence reveals relative fitness of bacteria in symbiotic and free-living environments. Proceedings of the National Academy of Sciences, 115:2425–2430.

Chaney, L. and R. S. Baucom. 2020. The soil microbial community alters patterns of selection on flowering time and fitness-related traits in Ipomoea purpurea. American Journal of Botany, 107:186–194.

Davis, M. R., F.-J. Zhao, and S. P. McGrath. 2004. Pollution-induced community tolerance of soil microbes in response to a zinc gradient. Environmental Toxicology and Chemistry: An International Journal, 23:2665–2672.

De Visser, J. A. G., T. F. Cooper, and S. F. Elena. 2011. The causes of epistasis. Proceedings of the Royal Society B: Biological Sciences, 278:3617–3624.

Dettman, J. R., C. Sirjusingh, L. M. Kohn, and J. B. Anderson. 2007. Incipient speciation by divergent adaptation and antagonistic epistasis in yeast. Nature, 447:585–588.

Dirilgen, N. and Y. Inel. 1994. Effects of zinc and copper on growth and metal accumulation in duckweed, Lemna minor. Bulletin of environmental contamination and toxicology, 53:442–449.

Dittami, S. M., L. Duboscq-Bidot, M. Perennou, A. Gobet, E. Corre, C. Boyen, and T. Tonon. 2016. Host–microbe interactions as a driver of acclimation to salinity gradients in brown algal cultures. The ISME journal, 10:51–63.

Epstein, H. E., H. A. Smith, G. Torda, and M. J. van Oppen. 2019. Microbiome engineering: enhancing climate resilience in corals. Frontiers in Ecology and the Environment, 17:100–108.

Falconer, D. S. 1996. Introduction to quantitative genetics. Pearson Education India.

Finkel, O. M., I. Salas-González, G. Castrillo, J. M. Conway, T. F. Law, P. J. P. L. Teixeira, E. D. Wilson, C. R. Fitzpatrick, C. D. Jones, and J. L. Dangl. 2020. A single bacterial genus maintains root growth in a complex microbiome. Nature, 587:103–108.

Fitzpatrick, C. R., Z. Mustafa, and J. Viliunas. 2019. Soil microbes alter plant fitness under competition and drought. Journal of evolutionary biology, 32:438–450.

French, E., I. Kaplan, A. Iyer-Pascuzzi, C. H. Nakatsu, and L. Enders. 2021. Emerging strategies for precision microbiome management in diverse agroecosystems. Nature Plants, 7:256–267.

Friesen, M. L. 2012. Widespread fitness alignment in the legume–rhizobium symbiosis. New Phytologist, 194:1096–1111.

Friesen, M. L., S. S. Porter, S. C. Stark, E. J. von Wettberg, J. L. Sachs, and E. Martinez-Romero. 2011. Microbially mediated plant functional traits. Annual Review of Ecology, Evolution, and Systematics, 42:23–46.

Göbel, P., C. Dierkes, and W. Coldewey. 2007. Storm water runoff concentration matrix for urban areas. Journal of Contaminant Hydrology, 91:26–42.

Gorton, A. J., D. A. Moeller, and P. Tiffin. 2018. Little plant, big city: a test of adaptation to urban environments in common ragweed (Ambrosia artemisiifolia). Proceedings of the Royal Society B, 285:20180968.

Gould, A. L., V. Zhang, L. Lamberti, E. W. Jones, B. Obadia, N. Korasidis, A. Gavryushkin, J. M. Carlson, N. Beerenwinkel, and W. B. Ludington. 2018. Microbiome interactions shape host fitness. Proceedings of the National Academy of Sciences, 115:E11951–E11960.

Grieneisen, L., M. Dasari, T. J. Gould, J. R. Björk, J.-C. Grenier, V. Yotova, D. Jansen, N. Gottel, J. B. Gordon, N. H. Learn, et al. 2021. Gut microbiome heritability is nearly universal but environmentally contingent. Science, 373:181–186.

Hadfield, J. D. 2010. MCMC methods for multi-response generalized linear mixed models: the MCMCglmm R package. Journal of Statistical Software, 33:1–22. Version 2.22.1.

Henke, R., M. Eberius, and K.-J. Appenroth. 2011. Induction of frond abscission by metals and other toxic compounds in Lemna minor. Aquatic Toxicology, 101:261–265.

Henry, L. P., M. Bruijning, S. K. Forsberg, and J. F. Ayroles. 2021. The microbiome extends host evolutionary potential. Nature Communications, 12:1–13.

Hereford, J. 2009. A quantitative survey of local adaptation and fitness trade-offs. The American Naturalist, 173:579–588.

Ho, K. H. E. 2017. The effects of asexuality and selfing on genetic diversity, the efficacy of selection and species persistence. Ph.D. thesis, University of Toronto St. George.

Honegger, R. 1993. Developmental biology of lichens. New Phytologist, 125:659–677.

Houwenhuyse, S., R. Stoks, S. Mukherjee, and E. Decaestecker. 2021. Locally adapted gut microbiomes mediate host stress tolerance. The ISME Journal, pages 1–14.

Jack, C. N., R. H. Petipas, T. E. Cheeke, J. L. Rowland, and M. L. Friesen. 2021. Microbial inoculants: silver bullet or microbial jurassic park? Trends in Microbiology, 29:299–308.

Jackrel, S. L., J. W. Yang, K. C. Schmidt, and V. J. Denef. 2021. Host specificity of microbiome assembly and its fitness effects in phytoplankton. The ISME Journal, 15:774–788.

Jain, D., R. Kour, A. A. Bhojiya, R. H. Meena, A. Singh, S. R. Mohanty, D. Rajpurohit, and K. D. Ameta. 2020. Zinc tolerant plant growth promoting bacteria alleviates phytotoxic effects of zinc on maize through zinc immobilization. Scientific Reports, 10:1–13.

Janssen, S., D. McDonald, A. Gonzalez, J. A. Navas-Molina, L. Jiang, Z. Z. Xu, K. Winker, D. M. Kado, E. Orwoll, M. Manary, et al. 2018. Phylogenetic placement of exact amplicon sequences improves associations with clinical information. Msystems, 3:e00021–18.

Jayasri, M. and K. Suthindhiran. 2017. Effect of zinc and lead on the physiological and biochemical properties of aquatic plant Lemna minor: its potential role in phytoremediation. Applied Water Science, 7:1247–1253.

Johnson, M. T. J. and J. Munshi-South. 2017. Evolution of life in urban environments. Science, 358.

Kern, E. M. A. and R. B. Langerhans. 2018. Urbanization drives contemporary evolution in stream fish. Global Change Biology, 24:3791–3803.

Kettlewell, H. B. D. 1955. Selection experiments on industrial melanism in the Lepidoptera. Heredity, 9:323–342.

Khan, S., R. Hauptman, and L. Kelly. 2021. Engineering the microbiome to prevent adverse events: challenges and opportunities. Annual Review of Pharmacology and Toxicology, 61:159–179.

Klinger, C. R., J. A. Lau, and K. D. Heath. 2016. Ecological genomics of mutualism decline in nitrogen-fixing bacteria. Proceedings of the Royal Society B: Biological Sciences, 283:20152563.

Krazčič, B., M. Slekovec-Golob, and J. Nemec. 1995. Promotion of flowering by Mn-EDDHA in the photoperiodically neutral plant Spirodela polyrrhiza (L.) Schleiden. Journal of Plant Physiology, 147:397–400.

Lau, J. A. and J. T. Lennon. 2012. Rapid responses of soil microorganisms improve plant fitness in novel environments. Proceedings of the National Academy of Sciences, 109:14058–14062.

Lebeis, S. L., S. H. Paredes, D. S. Lundberg, N. Breakfield, J. Gehring, M. McDonald, S. Malfatti, T. G. Del Rio, C. D. Jones, S. G. Tringe, et al. 2015. Salicylic acid modulates colonization of the root microbiome by specific bacterial taxa. Science, 349:860–864.

Li, E., R. de Jonge, C. Liu, H. Jiang, V.-P. Friman, C. M. Pieterse, P. A. Bakker, and A. Jousset. 2021. Rapid evolution of bacterial mutualism in the plant rhizosphere. Nature Communications, 12:1–13.

Liskco, Z. and J. Struger. 1996. Trace metals contamination of urban streams and stormwater detention ponds. In W. James, editor, Advances in modeling the management of stormwater impacts, chapter 17, pages 269–278. CRC Press.

Lozupone, C. and R. Knight. 2005. Unifrac: a new phylogenetic method for comparing microbial communities. Applied and environmental microbiology, 71:8228–8235.

Masoner, J. R., D. W. Kolpin, I. M. Cozzarelli, L. B. Barber, D. S. Burden, W. T. Foreman, K. J. Forshay, E. T. Furlong, J. F. Groves, M. L. Hladik, et al. 2019. Urban stormwater: An overlooked pathway of extensive mixed contaminants to surface and groundwaters in the United States. Environmental Science & Technology, 53:10070–10081.

McDonald, D., M. N. Price, J. Goodrich, E. P. Nawrocki, T. Z. DeSantis, A. Probst, G. L. Andersen, et al. 2012. An improved Greengenes taxonomy with explicit ranks for ecological and evolutionary analyses of bacteria and archaea. The ISME Journal, 6:610.

Morton, J. T., J. Sanders, R. A. Quinn, D. McDonald, A. Gonzalez, Y. Vázquez-Baeza, J. A. Navas-Molina, S. J. Song, J. L. Metcalf, E. R. Hyde, et al. 2017. Balance trees reveal microbial niche differentiation. MSystems, 2:e00162–16.

Newton, R. 1977. Abscisic acid effects on fronds and roots of Lemna minor L. American Journal of Botany, 64:45–49.

O’Brien, A. M., R. J. Sawers, S. Y. Strauss, and J. Ross-Ibarra. 2019. Adaptive phenotypic divergence in an annual grass differs across biotic contexts. Evolution, 73:2230–2246.

O’Brien, A. M., Z. H. Yu, D.-y. Luo, J. Laurich, E. Passeport, and M. E. Frederickson. 2019. Resilience to multiple stressors in an aquatic plant and its microbiome. American Journal of Botany.

Ontario Ministry of the Environment. 2011. Technical memorandum: an analysis of nutrients and select metals within wastewater (pond discharges). Technical report, Southwestern Regional Office.

O’Brien, A. M., C. N. Jack, M. L. Friesen, and M. E. Frederickson. 2021. Whose trait is it anyways? Coevolution of joint phenotypes and genetic architecture in mutualisms. Proceedings of the Royal Society B, 288:20202483.

O’Brien, A. M., J. Laurich, E. Lash, and M. E. Frederickson. 2020. Mutualistic outcomes across plant populations, microbes, and environments in the duckweed Lemna minor. Microbial Ecology, 80:384–397.

O’Brien, A. M., R. J. Sawers, J. Ross-Ibarra, and S. Y. Strauss. 2018. Evolutionary responses to conditionality in species interactions across environmental gradients. The American Naturalist, 192:715–730.

Panke-Buisse, K., S. Lee, and J. Kao-Kniffin. 2017. Cultivated sub-populations of soil microbiomes retain early flowering plant trait. Microbial Ecology, 73:394–403.

Panke-Buisse, K., A. C. Poole, J. K. Goodrich, R. E. Ley, and J. Kao-Kniffin. 2015. Selection on soil microbiomes reveals reproducible impacts on plant function. The ISME Journal, 9:980–989.

Paolacci, S., S. Harrison, and M. A. Jansen. 2018. The invasive duckweed Lemna minuta Kunth displays a different light utilisation strategy than native Lemna minor Linnaeus. Aquatic Botany, 146:8–14.

Peiffer, J. A., A. Spor, O. Koren, Z. Jin, S. G. Tringe, J. L. Dangl, E. S. Buckler, and R. E. Ley. 2013. Diversity and heritability of the maize rhizosphere microbiome under field conditions. Proceedings of the National Academy of Sciences, 110:6548–6553.

R Core Team. 2019. R: A Language and Environment for Statistical Computing. R Foundation for Statistical Computing, Vienna, Austria. Version 3.6.0.

Radić, S., M. Babić, D. Škobić, V. Roje, and B. Pevalek-Kozlina. 2010. Ecotoxicological effects of aluminum and zinc on growth and antioxidants in Lemna minor L. Ecotoxicology and Environmental Safety, 73:336–342.

RDSC. 2016. Rutgers duckweed stock center. Online; https://ruduckweed.org; accessed 25-October-2016.

Rebolleda-Gómez, M., N. J. Forrester, A. L. Russell, N. Wei, A. M. Fetters, J. D. Stephens, and T.-L. Ashman. 2019. Gazing into the anthosphere: considering how microbes influence floral evolution. New Phytologist, 224:1012–1020.

Rúa, M. A., A. Antoninka, P. M. Antunes, V. B. Chaudhary, C. Gehring, L. J. Lamit, B. J. Piculell, J. D. Bever, C. Zabinski, J. F. Meadow, M. J. Lajeunesse, B. G. Milligan, J. K. Karst, and J. D. Hoeksema. 2016. Home-field advantage? Evidence of local adaptation among plants, soil, and arbuscular mycorrhizal fungi through meta-analysis. BMC Evolutionary Biololgy, 16.

Sachs, J. L., U. G. Mueller, T. P. Wilcox, and J. J. Bull. 2004. The evolution of cooperation. The Quarterly Review of Biology, 79:135–160.

Santangelo, J. S., L. R. Rivkin, C. Advenard, and K. A. Thompson. 2020. Multivariate phenotypic divergence along an urbanization gradient. Biology Letters, 16:20200511.

Schneider, C. A., W. S. Rasband, and K. W. Eliceiri. 2012. NIH Image to ImageJ: 25 years of image analysis. Nature Methods, 9:671–675.

Schvartzman, M. S., M. Corso, N. Fataftah, M. Scheepers, C. Nouet, B. Bosman, M. Carnol, P. Motte, N. Verbruggen, and M. Hanikenne. 2018. Adaptation to high zinc depends on distinct mechanisms in metallicolous populations of Arabidopsis halleri. New Phytologist, 218:269–282.

Shantz, A. A., N. P. Lemoine, and D. E. Burkepile. 2016. Nutrient loading alters the performance of key nutrient exchange mutualisms. Ecology Letters, 19:20–28.

Spiegelhalter, D. J., N. G. Best, B. P. Carlin, and A. Van Der Linde. 2002. Bayesian measures of model complexity and fit. Journal of the Royal Statistical Society: Series B (Statistical Methodology), 64:583–639.

Thijs, S., M. Op De Beeck, B. Beckers, S. Truyens, V. Stevens, J. D. Van Hamme, N. Weyens, and J. Vangronsveld. 2017. Comparative evaluation of four bacteria-specific primer pairs for 16s rRNA gene surveys. Frontiers in Microbiology, 8:494.

Thrash, J. C. 2021. Towards culturing the microbe of your choice. Environmental Microbiology Reports, 13:36–41.

Tso, G. H. W., J. A. Reales-Calderon, A. S. M. Tan, X. Sem, G. T. T. Le, T. G. Tan, G. C. Lai, K. Srinivasan, M. Yurieva, W. Liao, et al. 2018. Experimental evolution of a fungal pathogen into a gut symbiont. Science, 362:589–595.

Turnbaugh, P. J., F. Bäckhed, L. Fulton, and J. I. Gordon. 2008. Diet-induced obesity is linked to marked but reversible alterations in the mouse distal gut microbiome. Cell Host & Microbe, 3:213–223.

Volkmer, B. and M. Heinemann. 2011. Condition-dependent cell volume and concentration of Escherichia coli to facilitate data conversion for systems biology modeling. PloS one, 6:e23126.

Wagner, M. R., D. S. Lundberg, D. Coleman-Derr, S. G. Tringe, J. L. Dangl, and T. Mitchell-Olds. 2014. Natural soil microbes alter flowering phenology and the intensity of selection on flowering time in a wild Arabidopsis relative. Ecology Letters, 17:717–726.

Wang, Z., D. Tsementzi, T. C. Williams, D. L. Juarez, S. K. Blinebry, N. S. Garcia, B. K. Sienkiewicz, K. T. Konstantinidis, Z. I. Johnson, and D. E. Hunt. 2021. Environmental stability impacts the differential sensitivity of marine microbiomes to increases in temperature and acidity. The ISME Journal, 15:19–28.

Weese, D. J., K. D. Heath, B. Dentinger, and J. A. Lau. 2015. Long-term nitrogen addition causes the evolution of less-cooperative mutualists. Evolution, 69:631–642.

Wippel, K., K. Tao, Y. Niu, R. Zgadzaj, N. Kiel, R. Guan, E. Dahms, P. Zhang, D. B. Jensen, E. Logemann, et al. 2021. Host preference and invasiveness of commensal bacteria in the Lotus and Arabidopsis root microbiota. Nature Microbiology, 6:1150–1162.

Wood, C. W. and E. D. Brodie III. 2015. Environmental effects on the structure of the G-matrix. Evolution, 69:2927–2940.

Wood, C. W. and E. D. Brodie III. 2016. Evolutionary response when selection and genetic variation covary across environments. Ecology Letters, 19:1189–1200.

Zahn, G. and A. S. Amend. 2019. Foliar fungi alter reproductive timing and allocation in Arabidopsis under normal and water-stressed conditions. Fungal Ecology, 41:101–106.

Zhang, M., Z. He, D. V. Calvert, P. J. Stoffella, and X. Yang. 2003. Surface runoff losses of copper and zinc in sandy soils. Journal of Environmental Quality, 32:909–915.

